# Interaction mapping of endoplasmic reticulum ubiquitin ligases identifies modulators of innate immune signalling

**DOI:** 10.1101/2020.03.18.993998

**Authors:** Emma J. Fenech, Federica Lari, Philip D. Charles, Roman Fischer, Marie Laétitia-Thézénas, Katrin Bagola, Adrianne W. Paton, James C. Paton, Mads Gyrd-Hansen, Benedikt M. Kessler, John C. Christianson

## Abstract

Ubiquitin ligases (E3s) embedded in the endoplasmic reticulum (ER) membrane regulate essential cellular activities including protein quality control, calcium flux, and sterol homeostasis. At least 25 different, transmembrane domain (TMD)-containing E3s are predicted to be ER-localised, but for most their organisation and cellular roles remain poorly defined. Using a comparative proteomic workflow, we mapped over 450 protein-protein interactions for 21 different stably expressed, full-length E3s. Bioinformatic analysis linked ER-E3s and their interactors to multiple homeostatic, regulatory, and metabolic pathways. Among these were four membrane-embedded interactors of RNF26, a polytopic E3 whose abundance is auto-regulated by ubiquitin-proteasome dependent degradation. RNF26 co-assembles with TMEM43, ENDOD1, TMEM33 and TMED1 to form a complex capable of modulating innate immune signalling through the cGAS-STING pathway. This RNF26 complex represents a new modulatory axis of STING and innate immune signalling at the ER membrane. Collectively, these data reveal the broad scope of regulation and differential functionalities mediated by ER-E3s for both membrane-tethered and cytoplasmic processes.

## Introduction

The endoplasmic reticulum (ER) is the largest membrane-bound organelle in eukaryotic cells, comprised of a complex network of sheets, tubules, junctions and contact sites that can occupy more than 35% of the entire cell volume^1^ and a significant fraction of the total membrane surface area. The continuous lattice it forms with the nuclear envelope (NE) makes extensive and dynamic contacts through distal projections with mitochondria^2^, peroxisomes^3^, endosomes^4,5^, plasma membrane^6^ and lipid droplets (reviewed in^7^). Within this extensive network, the ER accommodates biogenic, metabolic and regulatory multi-subunit transmembrane domain (TMD)-containing protein complexes that span the lipid bilayer and simultaneously carry out processes essential for cellular homeostasis.

Post-translational modification by ubiquitin (Ub) targets proteins for degradation, promotes interactions, directs subcellular localisation, or drives signalling^8^. The enzymatic cascade conjugating and extending Ub chains on proteins throughout the mammalian cell uses one (or more) of the > 600 Ub ligases (E3s) to provide reaction specificity by bringing substrates and Ub conjugating enzymes (E2s) in proximity^9^. E3s distinguish substrates either directly through dedicated binding domains/surfaces, or indirectly by assembling co-factors into specialised multi-subunit complexes with recognition and recruitment capabilities (reviewed in^10^). Within aqueous environments of the cytoplasm and nucleus, freely diffusing E3s access substrates with reduced spatial impediment. In contrast, E3s embedded within lipid bilayers by virtue of one or more TMDs or lipid anchor, have pre-determined orientation and lateral motion restricted to the planar membrane where they reside. Eukaryotic membrane-embedded E3s are found in the inner nuclear membrane (INM)^11^, ER^12^, mitochondria^9^, Golgi^13^, endosomes and plasma membrane^14^. The RING and HECT domains of E3s coordinating Ub transfer are exclusively exposed to the cytoplasm (and nucleus), enabling the access of cytosolic, nuclear, and proximal membrane proteins. Moreover, lumenal proteins from within the ER also reach the RING domain of at least one membrane-bound E3 (Hrd1), by way of an aqueous channel it forms within the lipid bilayer^15^. Thus, membrane bound E3s arguably serve as broad platforms for ubiquitination within the cell. Understanding how E3 complexes recognise substrates in and around membranes, how they can coordinate efficient Ub conjugation, how they regulate access to them, and which cellular processes they regulate, are important biological questions that remain outstanding.

ER-associated degradation (ERAD) has been the principal modality for understanding E3 function in the ER. Secretory cargo transiently or terminally misfolded during biogenesis in the ER is prone to aggregation that can cause proteotoxic stress, necessitating removal from the organelle that ultimately ends in its degradation^12,16,17^. ERAD serves in tandem with chaperone-mediated folding and assembly processes, acting as an integral facet of the organelle’s quality control machinery^18^. How we envisage E3 function at the ER has been shaped by extensive studies on the evolutionarily conserved Hrd1^19-26^. With as many as eight TMDs^15^, a cytoplasmic RING domain and an extended C-terminus with low complexity^27^, Hrd1 scaffolds specialised lumenal, integral membrane, and cytosolic co-factors such as the AAA-ATPase VCP/p97^28,29^ to form multi-component complexes that coordinate recognition, retrotranslocation, and ubiquitination of misfolded secretory cargo^20,24^.

E3 complexes not only control the quality of secretory cargo but also adjust the abundance (and hence activity) of ER-resident membrane proteins through “regulatory ERAD”^30^. As the primary site for phospholipid and sterol biosynthesis^31^, degradation of ER-resident rate-limiting enzymes such as HMGCoA reductase (HMGCR) by gp78/AMFR^32^, RNF139/Trc8^33^ and RNF145^34,35^, and squalene monooxygenase (SM) by MARCH6/Doa10^36,37^, help tune the output of this pathway to maintain homeostasis. Both gp78 and RNF145 use Insig1/2 as adaptors to recruit and degrade HMGCR^32,34^. Other processes at the ER membrane regulated by E3s and their interactors/adaptors include calcium flux^38^, innate immune signalling^39^, antigen presentation^40^, endosome diffusion^41^, ER morphology^42^ and apoptotic signalling^43^. E3 complexes represent important post-translational regulatory modules tending to the protein landscape of the mammalian ER, but most have not been extensively characterised. We developed a comparative proteomic workflow to define interactions of ER-resident E3s that sought to identify networks involved in maintaining ER and/or cellular homeostasis. Among the E3 interactors confidently identified were proteins previously reported in connection with lipid regulation, calcium flux, quality control, as well as new interactors involved in innate immune signalling.

## Results

### Isolation and discovery of ER-resident E3 interactors

Of the > 600 E3s present in the human proteome, about 10% contain TMDs that tether affiliated processes to lipid bilayers. Starting from previous reports^44-46^ and topology predictions, we shortlisted 25 E3s demonstrated (or predicted) to reside in the ER membrane. Selected ER-resident E3s (ER-E3s) are topologically and structurally disparate, arise from different phylogenetic lineages, but commonly possess one or more TMDs and a cytosolic RING-domain (Fig. 1a). With a design to determine cognate ER-E3 co-factors and substrates, we developed a generalised expression protocol coupled to a comparative immunoprecipitation liquid chromatography tandem mass spectrometry (IP-LC-MS/MS) workflow that enriched for proteins interacting at the ER. We generated 25 stable, individual, HEK293 cell lines, each expressing a FLAG-HA (FH)-tagged E3 inducible by doxycycline (DOX) using Flp-In^™^ recombination (see Methods, Fig. 1b). Positioning of the FH-tag at either E3 terminus was determined by taking into account prior experimental observations^44^, predicted topology, and proximity to RING domains or other prominent structural features (Fig. 1a, Extended Data Table 1). Variable induction parameters sought to produce comparable E3 expression levels from each cell line (Extended Data Fig. 1a). More than 80% of ER-E3s tested (21/25) could be detected from whole cell lysates (WCL), accumulating in a DOX-dependent manner that reflects relative stability (Extended Data Fig. 1a). Including MG132 along with DOX enhanced detection of some E3s (e.g. RNF26, BFAR, TMEM129), indicative of intrinsic instability and constitutive turnover by proteasome-mediated degradation. E3s colocalised exclusively (or partly) with the ER-markers calnexin or KDEL (Extended Data Fig. 1b), consistent with their expected residency in the ER.

**Fig. 1.**
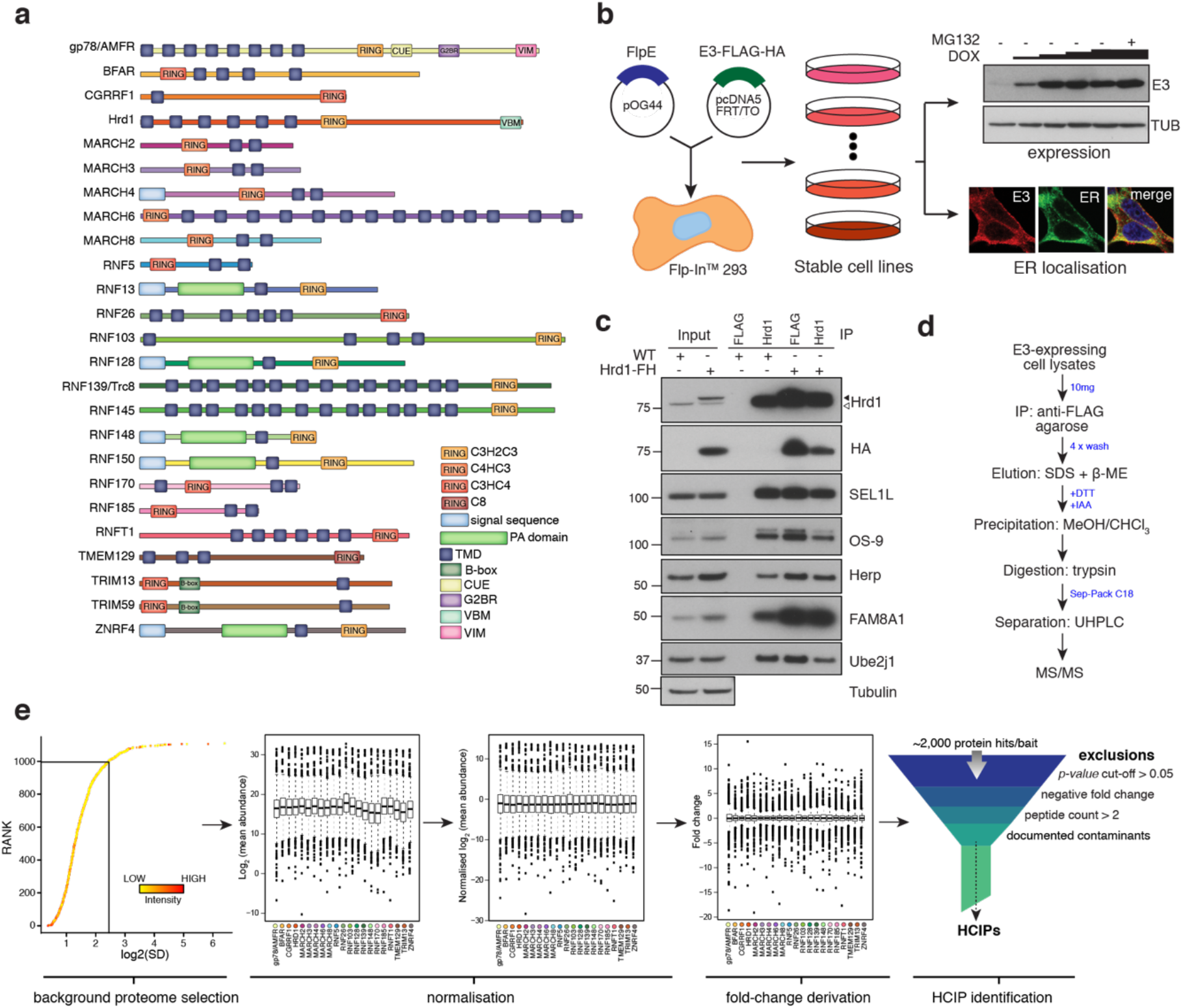
Proteomic analysis of ER-resident ubiquitin ligases. (**a**) ER-resident E3s and their predicted domains. (**b**) Workflow to generate and validate Flp-In^™^293 cell lines stably expressing FLAG-HA-tagged E3s (FH-E3 or E3-FH). Each Flp-In^™^293 cell line stably integrating a tagged E3 was screened for induction and expression over increasing concentrations of DOX and MG132 treatment by western blot (anti-FLAG) as well residency in the ER by immunofluorescence, evaluating colocalisation with markers of the ER, calnexin or KDEL. (**c**) Coimmunoprecipitation profiles of endogenous Hrd1 and DOX-induced Hrd1-FH prepared in 1% LMNG and isolated by anti-Hrd1 or anti-FLAG, as indicated. Input (20% of total IP) is also shown. (**d**) Workflow of sample preparation for LC-MS/MS analysis. (**e**) Bioinformatic processing pipeline for identification of high-confidence candidate interacting proteins (HCIPs) for the E3 baits.

To assess whether tagged ER-E3s faithfully reproduce endogenous complexes, we compared the interaction profile of Hrd1^24,26,27,47-49^ with that of DOX-induced Hrd1-FH. Cofactors including SEL1L, FAM8A1, OS-9, Herp, and UBE2J1 were comparably coimmunoprecipitated (co-IPed) by both Hrd1 and Hrd1-FH (Fig. 1c). Moreover, velocity sedimentation revealed that Hrd1-FH complexes migrate in fractions that overlap with those formed by endogenous Hrd1 (Extended Data Fig. 1c). From this, we anticipate that *bona fide* protein-protein interactions of candidate E3s will be recapitulated by the exogenously expressed E3s.

Preserving native interactions between TMD-containing E3s and co-factors (or substrates) during sample processing was essential to ensure robust detection by liquid chromatography tandem mass spectrometry (LC-MS/MS). Cells were solubilised using LMNG (Lauryl Mannose Neopentyl Glycol)-containing buffer, a detergent shown previously to preserve labile ER-E3 complex interactions^27^. Immunoprecipitated E3-interactor complexes were washed and subsequently eluted from beads non-selectively by SDS to obtain the sample complexity necessary for subsequent comparative analyses (see below). All samples were prepared and processed for LC-MS/MS in parallel to facilitate comparative analysis (Fig. 1d, see details in Materials and Methods). ~1,600 individual protein groups were detected within each sample. In total, > 2,000 unique proteins were identified (Extended Data Table 2). To distinguish each E3’s most relevant interactors, we adapted the Bait-Specific Control Group (BSCG) method described previously for multiple sample processing^50^ (Fig. 1e, Extended Data Methods). This method defines the set of commonly detected interactors common as the “background proteome” and normalises each sample to it to facilitate multi-sample comparison. By permitting relative fold-change and p-values to be determined for each identified protein, normalisation enables an enrichment to be assessed. To enrich for factors with relevant change, each putative E3 interactor was individually evaluated for: (1) p-value <0.05; (2) positive fold-change; and (3) >1 unique peptide (Fig. 1d). Commonly identified contaminants (Extended Data Table 3) were excluded and removed from the dataset. Proteins meeting all criteria were designated as “high-confidence candidate interacting proteins” (HCIPs)^51^.

An inherent limitation of BSCG analysis is that candidate interactors detected in only one E3 sample evade classification as HCIPs because a relative fold-change cannot be calculated. Since exclusive ER-E3 interactors represent high-value candidates, we also analysed raw data by calculating the semi-quantitative spectral index quantitation (SINQ) score for each sample (as described in^52^). Filtering for sample-exclusive, high SINQ score proteins in each sample identified an additional 28 interactors co-precipitating uniquely by specific ER-E3 (Extended Data Table 4). Merging the modified BSCG and SINQ analyses revealed over 428 interactions with 21 E3s (Extended Data Fig. 1d), the majority of which are unreported. This composite dataset represents a systematic attempt to define E3 complexes at the ER membrane.

### Interaction landscape of ER-resident E3s

From the 21 E3 ligases we comparatively examined, 218 different HCIPs that formed 400 interactions were identified (Extended Data Table 5). Hierarchical clustering of individual E3 interactomes (Fig. 2a) revealed that HCIPs were both exclusive to and shared between ER-E3s. Visualising HCIP networks for E3s individually (Extended Data Fig. 2) revealed interactors brought down with varying degrees of confidence and linked to a diverse range of activities, as exemplified by the networks for both Hrd1 (Fig. 2b) and RNF185 (Fig. 2c). While ER-E3 raw abundance differed markedly, this did not correlate with the number of HCIPs identified (Fig. 2d). Therefore, the number of interactions detected was a function of intrinsic E3 properties and not simply expression level. Importantly, over 50% of HCIPs (112/218) were significantly enriched by just one ER-E3 while those remaining associated with 2 (67/218), 3 (19/218) or > 4 (20/218) different ligases (Fig. 2e). HCIPs enriched by only one ER-E3 might represent specific cofactors or cognate substrates whereas interaction with multiple E3 could reflect proteins with generalised or adaptable functionality. We excluded HCIPs identified for RNF145, RNF150, Trim59 and RNF13 because bait peptide counts were insufficient (<2) or interactors failed to be enriched.

**Fig. 2.**
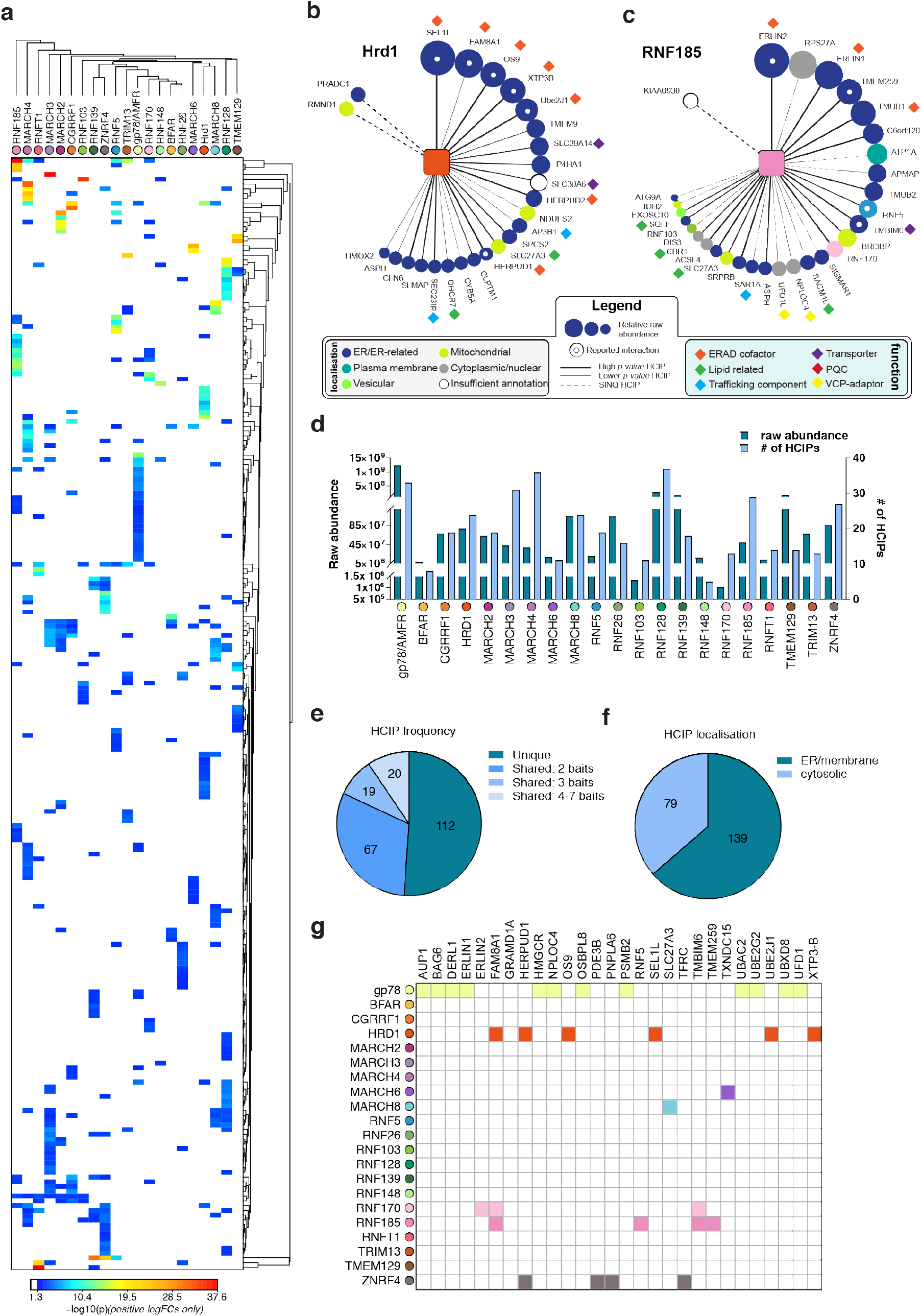
Interaction landscape of 21 ER-resident E3s. (**a**) Hierarchical clustering of the 21 E3s and their associated HCIPs represented as a heat map, where the colours of individual interactors correspond to their calculated p-values. Representative HCIP interaction wheels for (**b**) Hrd1 and (**c**) RNF185. Parameters represented are described in the adjoining legend. (**d**) Raw abundance (RA) and number of HCIPs determined for each E3. (**e**) Distribution of HCIP interactions with E3s as unique or shared. (**f**) Classification of HCIPs as ER/membrane or cytosolic proteins as defined by presence of validated and predicted signal peptides, glycosylation sites, disulphide bonds, and transmembrane domains (UniProt). (**g**) E3-HCIPs interactions identified previously in the BioGRID database (https://thebiogrid.org/).

To concentrate subsequent analyses on interactions made at the ER, we searched the HCIP dataset for predicted protein features/domains associated with organelle targeting or residency, such as TMDs, signal sequences, N-linked glycosylation or disulphide bonds (UniProt). Approximately two-thirds of HCIPs (139/218) contained features consistent with localisation to the endomembrane system, validating our enrichment strategy for ER-associated proteins (Fig. 2f). Originality of E3-HCIP interactions is reflected by their underrepresentation in current protein-protein interaction resources (e.g. BioGRID 3.0), where only ~7% of E3-HCIP interactions have been reported previously^53^ (Fig. 2g). Unsurprisingly, most of these reported interactions (18/31) were with either of the two extensively characterised ER-E3s, Hrd1 and gp78/AMFR. This consistency provides additional assurance that the workflow could identify bona fide ER-E3 interactors.

### ER-E3s, ERAD, and ER stress

E3 functionality at the ER is most commonly associated with the quality control (QC) process of ERAD (reviewed in^12,17^). We investigated whether any ER-E3s enriched for HCIPs implicated previously in ERAD, such as those that comprise the Hrd1 complex (e.g. SEL1L, UBE2J1, OS-9)^24,26^. While all were associated with Hrd1 (Fig. 2b), they were not HCIPs prominently enriched by other ER-E3s (Fig. 3a). Established gp78/AMFR interactors including UBE2G2, Derlin1/DERL1, and UBAC2^24,54^, also did not feature among HCIPs of other E3s (Extended Data Fig. 2a, Fig. 3a). Thus, factors previously linked with ERAD do not appear to have generalised functionality that is readily adopted by other ER-E3s, except for VCP/p97 (discussed below).

**Fig. 3.**
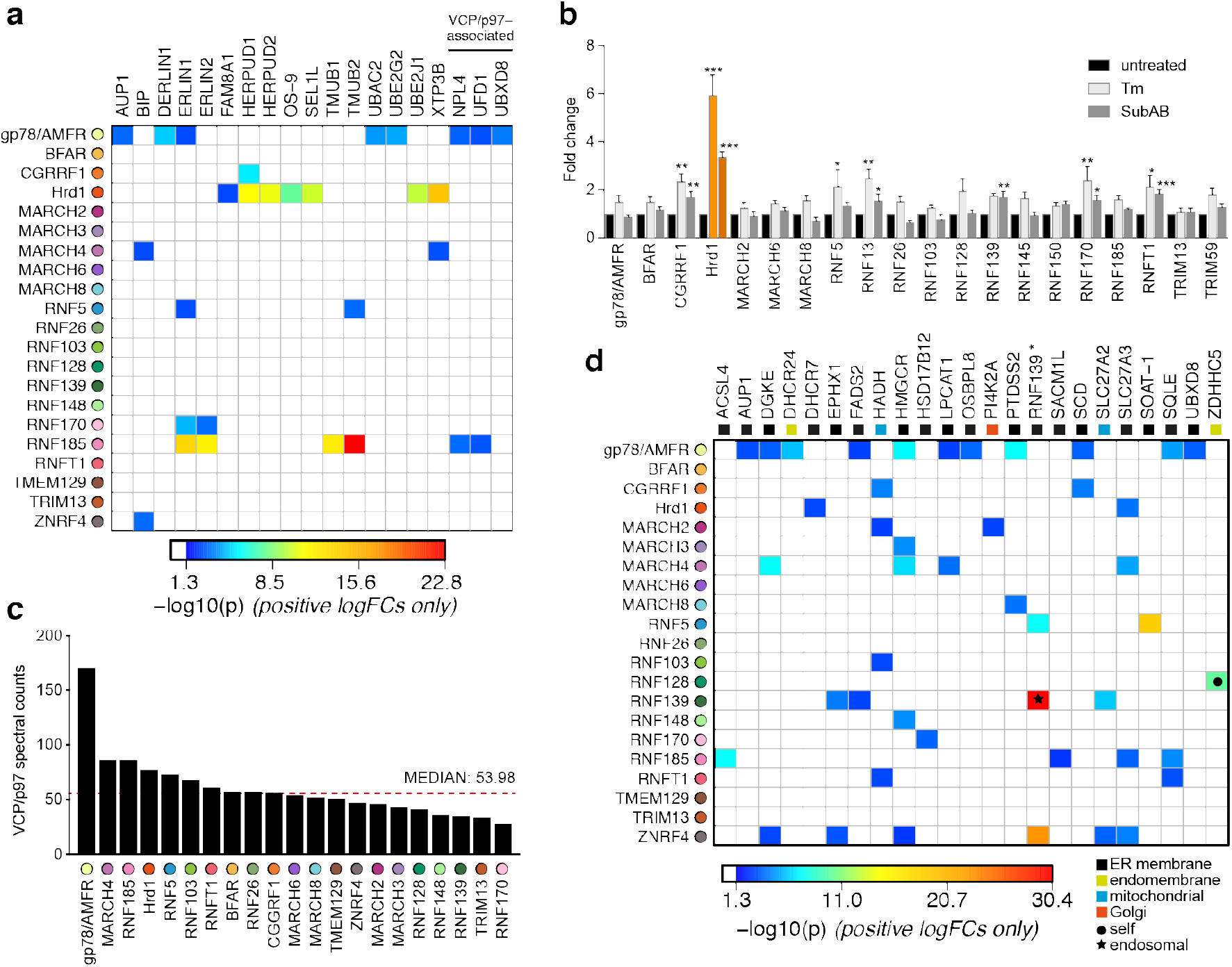
Functional associations of E3 and HCIPs. (**a**) Heatmap depicting established ERAD components found as HCIPs with the panel of ER-resident E3s, with colours of individual interactors corresponding to their calculated p-values. (**b**) Transcriptional analysis of parental Flp-In^™^293 cells determined by NanoString. Data depict fold change of E3 transcripts measured from tunicamycin-treated (Tm, 500 ng/ml, 8 h) and SubAB-treated cells when compared to untreated. Mean and S.E.M. are shown from three biological repeats (n=3). *\*P* < 0.05, \*\**P* < 0.01, *\*\*\*P* < 0.001. Detailed statistical analysis can be found in Extended Data Table 12. Hrd1 is highlighted in orange for reference. (**c**) Absolute number of spectral counts detected for VCP/p97 for each ER-resident E3 determined by SINQ analysis. The dotted red line shows the median spectral counts for reference (**d**) Heatmap representing the association between proteins involved in lipid regulation (synthesis; metabolism and transport) and E3 baits. Colours associated with individual interactors correspond to their calculated p-values.

One way the unfolded protein response (UPR) resolves ER stress is by coordinated upregulation of Hrd1 (and its cofactors) to increase ERAD capacity^55^. We investigated whether other ER-E3s respond similarly to ER stress by quantitatively monitoring transcriptional changes in ER-E3s and factors linked to ERQC in HEK293 cells treated with Tunicamycin (Tm) or the AB5 family bacterial toxin Subtilase cytotoxin (SubAB5) toxin^56^ (Extended Data Table 6). CGRRF1, RNF13, RNF170 and RNFT1 transcript levels increased with acute ER stress (~2 fold) as reported previously^57^, along with RNF5 (Tm only) and RNF139 (SubAB5 only) (Fig. 3b). When compared to the ~6-fold change (for Tm) observed for Hrd1, however, any responsive contribution made by other E3s to ER stress resolution may be nominal. Of the 226 HCIPs identified, 24 are among the 278 targets of the UPR transcription factors XBP1, ATF6, and ATF4 collated previously^58^, with a quarter (6/24) represented by the Hrd1 complex alone (Extended Data Table 7). Moreover, ER homeostatic maintenance did not require any individual ER-E3 since siRNA-mediated knockdowns of endogenous isoforms were not sufficient to induce the splicing of *XBP1* (Extended Data Fig. 3a), consistent with findings from CRISPRi screens for ER stress induction^59^. Taken together, these findings are consistent with unique positioning of Hrd1 among E3s to resolve proteotoxic ER stress^60^.

### Recruitment of VCP/p97 to ER E3 complexes

Binding and hydrolysis of ATP enables VCP/p97 to generate the force necessary to extract polypeptides from the ER membrane during ERAD^29,61^. VCP/p97 enrichment could therefore reflect a need for substrate extraction and a role in ERAD, so we searched for reported ER-E3s and HCIPs interactions with the AAA ATPase. VCP/p97 is recruited to protein complexes throughout the cell by factors that contain a UBX (Ub-regulatory X) domain or by a linear sequences such as the SHP motif (also known as binding site 1, BS1), the VCP-interacting motif (VIM), or the VCP-binding motif (VBM)^62^. These domains/motifs can be found in ER-E3s^63,64^, their cofactors (e.g. Derlins, VIMP)^65,66^ or ERAD-related enzymes (e.g RHBDL4)^67^. Surprisingly, only two ER-related HCIPs containing confirmed VCP-binding domains were found. FAF2/UBXD8 containing a UBX domain^48^ and Derlin1 containing a SHP domain^65^, were both associated with gp78/AMFR (Extended Data Fig. 2a, Fig. 3a) as previously reported^24^.

Differentiating bona fide recruitment of VCP/p97 from non-specific binding can be problematic in an IP-LC-MS/MS workflow because it is highly abundant, involved in a broad range of cellular processes, and can interact non-specifically. To address this, we compared VCP/p97 spectral counts determined for each E3 IP to assess whether any enriched the AAA ATPase. Among the highest were those for gp78/AMFR and Hrd1, in line with their established recruitment for ERAD, but RNF185, MARCH4 and RNF5 were also among the upper quartile (Fig. 3c). The soluble VCP/p97 cofactors UFD1 and NPLOC4 were also HCIPs of RNF185 (discussed below), which lends additional support for this E3 in an ERAD-related role^68^. RNF185 was recently highlighted among the set of prominent candidates identified in the VCP/p97 interactome^69^. Despite this, neither RNF185 nor its HCIPs contained canonical VCP/p97 binding domains/motifs that might justify recruitment. An unidentified factor or the presence of a non-canonical (or cryptic) VCP/p97 binding motif^70^ could be responsible, but this remains undetermined.

### Ubiquitin-related HCIPs and ER-E3s

ER-E3 HCIPs included factors implicated in Ub conjugation and binding, as well as those containing a Ub-like (UBL) domain. However, other than the two well-characterised E2-E3 pairs UBE2G2-gp78/AMFR and UBE2J1-Hrd1, our isolation protocol does not appear to have robustly preserved these interactions. As almost all E2s are soluble and bind with moderate or weak affinity to E3 RING domains during Ub transfer, their scarcity was not unexpected. The Ub precursor, ribosomal fusion protein RSP27A, was enriched by both RNF185 and gp78/AMFR, reflecting either its binding to or direct modification of the E3s. Hrd1 and RNF185 enriched UBL domain-containing proteins (and homologues) Herp/Herp2 (HERPUD1/HERPUD2) and TMUB1/TMUB2, respectively (figs. 2c, 3a). Both have been linked to ERAD; TMUB1 links Erlin1 to gp78^71^ while Herp binds to FAM8A1^27^ and activates Hrd1 through a cytoplasmic domain^72^. Deubiquitinating enzymes (DUBs) were not among E3 HCIPs, which may not be surprising as the DUB interactome did not report interaction with any ER-E3s^51^. Some ER-E3s did co-precipitate (e.g. RNF185-RNF170, RNF185-RNF5), and may reflect organisation consistent with coordinated or sequential ubiquitination as part of ERAD^73,74^ or alternatively, an E3-substrate relationship.

### ER-E3s and calcium-related HCIPs

Signalling from G-protein-coupled receptors (GPCRs) activates inositol 1,4,5-triphosphate (IP3)-receptors (IP3Rs), causing this calcium-gated Ca^2+^ channel to be ubiquitinated and turned over from the ER membrane^75^ by RNF170 and its cofactors Erlin1/Erlin2^38,76^. We identified enrichment of the ERLIN1/2 heterodimer by RNF170 and RNF185, and Erlin1 alone by RNF5 and gp78 (Figs. 2c, 3a, Extended Data Fig. 2). RNF170 is an HCIP of both RNF185 and gp78, which suggests Erlin1/2 interactions could serve as a bridge for larger heterooligomeric E3 complexes. IP3R was only enriched by RNF170, in line with its previous identification as a cognate substrate and demonstrating that this methodology can also identify *bona fide* E3 substrates. Consistent with a larger hetero-oligomer, RNF170 and RNF185 share other HCIPs including the putative secreted factor c6orf120 and TMBIM6/BI-1/Bax-inhibitor 1 (Extended Data Fig. 3b). These form a Ca^2+^ leak channel in the ER that protects cells from ER stress^77^ by regulating Ca^2+^ release and interacting with TMBIM3/GRINA (see^78^). RNF185 enriched for TMUB1/TMUB2 and TMEM259/Membralin (Fig. 2c), a polytopic ER protein linked to motor neuron survival^79^ that appears to have been erroneously assigned as part of the Hrd1-gp78 ERAD network^80^. RNF185 interactions were validated by co-expression and pulldown with S-tagged HCIPs (Extended Data Fig. 3c). Interestingly, the Ca^2+^-load-activated Ca^2+^ channel TMCO1, which prevents overfilling in the ER^81^, was also enriched by RNF170. Collectively, RNF170 and RNF185 appear to associate with proteins linked to homeostatic maintenance of ER Ca^2+^ levels related to ER stress and apoptosis.

### E3s and lipid-related HCIPs

Sterols and fatty acids are produced through coordinated biosynthetic reactions at the ER membrane. 3-hydroxy-3-methylglutaryl-CoA-reductase (HMGCR) and squalene epoxidase (SQLE), are among the best examples of rate-limiting enzymes degraded by ER-E3s through negative feedback to regulate biosynthetic activity. 25 different HCIPs were involved in the biosynthesis and regulation of cholesterol, fatty acids, or phospholipids, with 80% annotated as TMD-containing proteins residing in the ER (Fig. 3d). Sterol and fatty acid-related HCIPs were enriched by 16 different E3s with nearly a quarter of all interactions (11/48) made with gp78/AMFR. Among the HCIPs of gp78/AMFR was HMGCR, previously reported as its substrate^33^. Conventionally, lipid-related HCIPs associating with E3s would do so as substrates for regulatory ERAD processes and may include DHCR7 (Hrd1), ACSL4 (RNF185) or SOAT1 (RNF5). Additionally, ZDHCC5 (RNF128) is a palmitoyl-acyltransferase which itself is palmitoylated^82,83^ and involved in endosome-Golgi trafficking^84^. Notably, it is one of the few palmitoyl transferases localised to the endosomal system, consistent with RNF128 (GRAIL) also reportedly present beyond the ER in endocytic compartments^85^.

### RNF26 is unstable and degraded by the UPS

ER-E3s co-precipitated uncharacterised proteins with high specificity and abundance, which may reflect essential functionality at the ER. One example is the HCIPs and complexes formed by RNF26, an ER-E3 implicated previously in innate immune signalling^86^ and organisation of perinuclear endosomes^41^. RNF26 is integrated into the ER membrane through six predicted TMDs (Fig. 4a). Its canonical C3-H-C4-type RING domain lying near its C-terminus is evolutionarily conserved (Extended Data Fig. 4a) and shares sequence and positional similarity with the nuclear SUMO-targeted Ub ligase (StUbL) RNF4^87,88^, Inhibitor of Apoptosis Proteins (IAPs,^89^), and MDM2^90^ (Fig. 4b), all of which form homo-/heterodimers through their RING domains. Despite efforts and consistent with published data^41^, endogenous RNF26 protein could not be detected from whole cell lysates, even though relative mRNA levels in HEK293 cells were comparable to other E3s such as Hrd1 (Extended Data Fig. 4b). Consequently, we used DOX-induced expression of FH-RNF26_WT_ to facilitate detection by immunoblotting, which was enhanced by treatment with MG132 (Fig. 4c, top panel, Extended Data Fig. 1a). Stabilisation by MG132 reflected intrinsic instability of RNF26 and suggested constitutive disposal by ERAD or an ERAD-like process. RNF4 conjugates Ub using a penultimate tyrosine (Y189) to engage the E2-Ub thioester^91^. This aromatic residue is conserved in RNF26 (Y432, Fig. 4a,b, Extended Data Fig. 4a), where a neutralising mutation (Y432A) stabilises FH-RNF26 protein levels and renders it insensitive to MG132 (Fig. 4c, middle panel). Radiolabel pulse-chase assays confirmed the increased stability of newly synthesised FH-RNF26_Y432A_ compared to FH-RNF26_WT_ (t_1/2_ ~ 2 hrs. vs. < 0.5 hrs, Fig. 4d). Expression of FH-RNF26_Y432A_ also enabled endogenous RNF26 to be co-precipitated and detected (Fig. 4e), consistent with the formation of stable and inactive heterodimers.

**Fig. 4.**
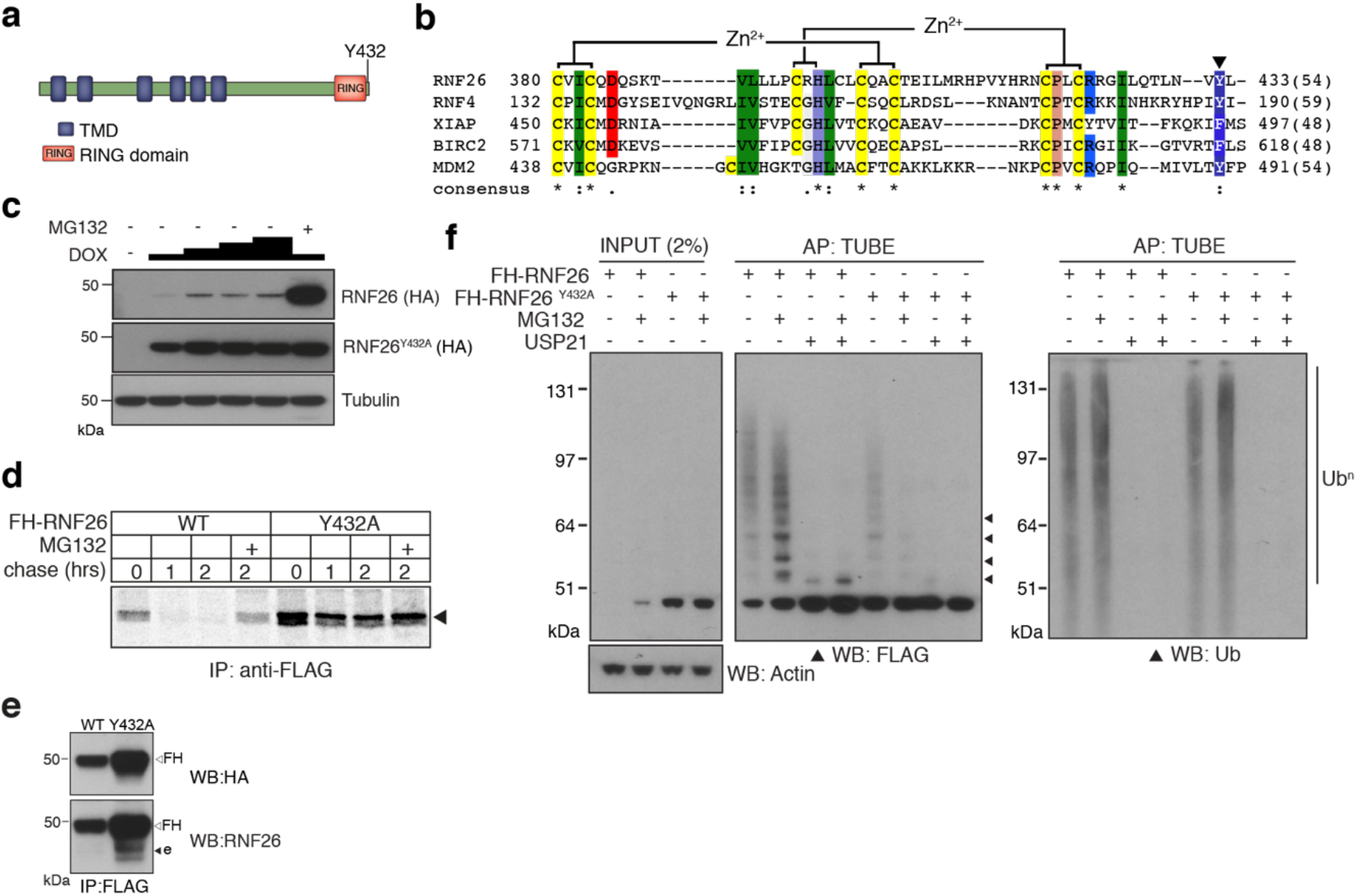
Characterisation of the RNF26 ubiquitin ligase complex. (**a**) Domain organisation of human RNF26 protein. (**b**) Protein sequence alignment of the RING domain and C-terminus of RNF26 with those of human RNF4 (P78317), XIAP (P98170), BIRC2 (Q13490) and MDM2 (Q00987). Conserved residues are demarcated according to the Rasmol colour scheme. (**c**) DOX titration of Flp-In^™^293 cells stably expressing FH-RNF26_WT_ or FH-RNF26_Y432A_ ± MG132 with lysates separated by SDS-PAGE and resulting western blots probed for RNF26 (anti-HA) and tubulin. (**d**)^35^S-Met/Cys pulse-chase assay (0, 1, 2 h) of DOX-induced Flp-In^™^293 cells stably expressing FH-RNF26_WT_ or FH-RNF26_Y432A_, ± MG132 and immunoprecipitated by anti-FLAG agarose. (**e**) Co-immunoprecipitation of endogenous RNF26 from DOX-induced Flp-In^™^293 cells stably expressing FH-RNF26_WT_ or FH-RNF26_Y432A_ by anti-FLAG beads. Detection of FLAG-HA (FH) and endogenous (e) RNF26 by western blot using the indicated antibodies. (**f**) TUBE pulldowns from FH-RNF26_WT_ or FH-RNF26_Y432A_ Flp-In^™^293 cell lysates, ± MG132 and Usp21.

Like other E3 dimers, the increased stability afforded by MG132 and the E2-Ub activating mutant (Y432A) is consistent with RNF26 auto-ubiquitination. Accordingly, immunoprecipitated FH-RNF26_WT_ produced FLAG-immunoreactive ladders and high-molecular weight smears, enhanced by MG132 and collapsed by the non-selective DUB Usp21 (Fig. 4f). *In vitro* ubiquitination assays using immunopurified FH-RNF26 (WT or Y432A) and recombinant UbcH5a, faithfully recapitulated these observations (Extended Data Fig. 4c). Curiously however, immunoprecipitated FH-RNF26_WT_ subjected to a panel of linkage specific DUBs indicated RNF26 modification by Ub chains not conventionally linked with degradation, including K33-, K63- and K29-linkages as well as multiple mono-ubiquitination (Extended Data Fig. 4d). Linkage diversity in RNF26 ubiquitination may indicate effects not only on turnover, but also on scaffolding and interaction with other proteins.

### Discovery of RNF26 interactors that enrich with stable mutant

The intrinsic instability of RNF26_WT_ may have led low abundance interactors to be below detection thresholds, and so IP-LC-MS/MS was also performed using FH-RNF26_Y432A_, which was then introduced into the BSCG and SINQ analyses (Fig. 5a). The inclusion of FH-RNF26_Y432A_ increased the total number of high-confidence protein-protein interactions from 428 to 457. A comparison of RNF26_WT_ and RNF26_Y432A_ HCIPs revealed many of the same interactors, albeit now with more specific enrichment (i.e. higher p-values) (Fig. 5b, Extended Data Table 8). We prioritized TMD-containing and ER-related HCIPs for validation, selecting first among those enriched by both forms. Prominent HCIPs of both RNF26 forms included TMEM43/LUMA (Fig. 5b, Extended Data Fig. 5a), an evolutionarily ancient multi-spanning membrane protein present in both the ER and INM^92,93^, and ENDOD1 (endonuclease domain containing-1), an uncharacterised polytopic protein containing a putative DNA/RNA nonspecific nuclease domain (Extended Data Fig. 5a). RNF26_Y432A_ also enriched for TMED1, an atypical member of the p24 cargo receptor family^94^, an ER membrane-shaping factor TMEM33^95^, as well as the ERAD and lipid droplet-related proteins UBXD8 and AUP1 (Fig. 5a,b, Extended Data Fig. 5a).

**Fig. 5.**
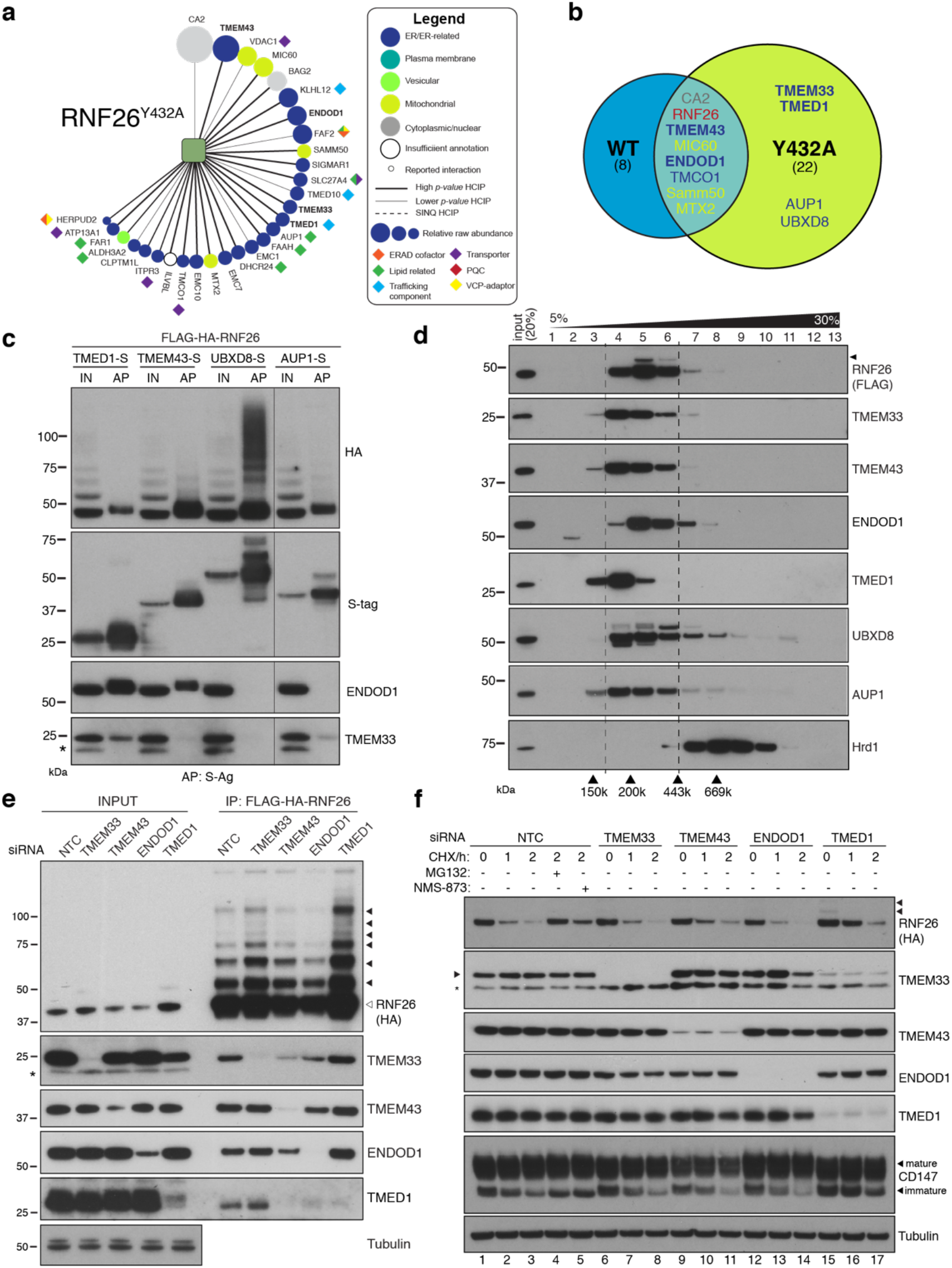
RNF26 assembles with HCIPs in the ER. (**a**) HCIP interaction network wheel for FH-RNF26_Y432A_. Legend described in Extended Data Fig. 2. (**b**) Venn diagram of HCIPs identified by LC-MS/MS for FH-RNF26_WT_ and FH-RNF26_Y432A_. ER-resident proteins are indicated in white. (**c**) Co-precipitation of FH-RNF26_WT_ from stable Flp-In^™^293 cells by transiently expressed S-tagged HCIPs (TMED1, TMEM43, UBXD8 and AUP1). Cells were solubilised in 1% LMNG and protein complexes affinity purified from the resulting lysates by S-protein agarose. Western blots of affinity purified material (AP) and input lysate (IN) were probed with antibodies recognising RNF26 (HA), HCIPs (S-tag), TMEM33 and ENDOD1. (**d**) Velocity sedimentation of FH-RNF26 complexes from 1% LMNG-solubilised lysates on a sucrose gradient (5-30%), with individual TCA-precipitated fractions (1-13) subsequently separated by SDS-PAGE and the resulting western blots probed for the indicated proteins. Ubiquitinated forms of RNF26 are indicated by black arrowheads. Molecular weight of gel filtration standards solubilised and sedimented in equivalent buffer conditions are shown for comparison beneath the Hrd1 blot, which in turn serves to highlight a complex of a different mass. (**e**) siRNA mediated knockdown of HCIPs in FH-RNF26_WT_ Flp-In^™^293 cells alters interaction profiles. RNF26 complexes immunoprecipitated by anti-FLAG agarose were separated by SDS-PAGE with resulting western blots probed by antibodies for RNF26 and the indicated HCIPs. Tubulin was used as a loading control. (**f**) Cycloheximide (CHX) chase assays (100 μg/ml; 0, 1, 2 h) of FH-RNF26_WT_ Flp-In^™^293 cells (DOX, 1μg/mL, 18 h) knocked down for individual HCIPs by siRNAs with the resulting western blots probed for the indicated antibodies. MG132 (10μM) and NMS-873 (10μM) were included with NTC samples where indicated. Ubiquitinated forms of RNF26 are denoted by black arrowheads. Mature and immature forms of CD147 are depicted by white arrowheads.

To validate interactions and gain insight into quaternary structures, S-tagged HCIPs were coexpressed with FH-RNF26_WT_ and the resulting co-precipitation profiles compared. FH-RNF26 was reproducibly brought down by S-tagged TMEM43, TMED1, AUP1 and UBXD8, with the latter exhibiting a high molecular weight smear of RNF26 reminiscent of polyubiquitinated forms (Fig. 5c). For reasons that are not clear, exogenous expression of S-tagged ENDOD1 and TMEM33 proved difficult to detect, however S-tagged TMEM43 and TMED1 were able to bring down endogenous ENDOD1 and TMEM33 (TMED1 only), indicating that these HCIPs comprised larger macromolecular complexes. Neither ENDOD1 nor TMEM33 were co-precipitated by S-tagged AUP1 or UBXD8, suggesting the formation of heterogeneous complexes by RNF26. Velocity sedimentation of lysates from FH-RNF26 expressing cells revealed a profile of a complex migrating between ~200-300kDa (fractions 4-6) with endogenous TMEM43, TMED1, ENDOD1, TMEM33, UBXD8 and AUP1 co-sedimenting with FH-RNF26, albeit with varying degrees of overlap within fractions (Fig. 5d). The robustness of RNF26-HCIP interactions was assessed by performing IPs in Triton X-100 (TX-100) as well as the milder LMNG; detergents which differ in their ability to stabilise ER membrane protein complexes (^27^, Extended Data Fig. 5b,c). TMEM43 co-precipitated by FH-RNF26_Y432A_ was independent of detergent conditions while other HCIP interactions were compromised to varying degrees in TX-100. These data support the formation of one or more heteromeric complexes containing RNF26 and HCIPs with a key interaction likely made through TMEM43.

To investigate organisation of RNF26 complexes in greater detail, validated siRNAs targeting HCIPs (Extended Data Figs. 5d-g) were introduced into FH-RNF26_WT_ expressing cells and co-precipitation profiles monitored. Silencing TMEM43 disrupted the ability of FH-RNF26_WT_ to coprecipitate other HCIPs, without substantially altering total cellular levels (Fig. 5e). A reduction in immunoprecipitated RNF26 (both unmodified and ubiquitinated) resulted from ENDOD1 knockdown, which might underlie the lower levels of other HCIP levels in RNF26 pulldowns, in particular TMED1. Although the pattern of HCIPs co-precipitated by RNF26 was unaffected by depletion of either TMEM33 or TMED1, detection of RNF26 and RNF26-Ub^n^ forms was markedly enhanced by the loss of TMED1. These profile changes, together with the fact that the RNF26-TMEM43 interaction is resistant to solubilisation in TX100, indicate TMEM43 plays a key role in RNF26 complex formation, while ENDOD1 and TMED1 exert influence over RNF26 abundance.

We next asked whether HCIPs affected RNF26 stability (Fig. 4e). In cycloheximide (CHX) chase assays, siRNA-mediated depletion of TMEM43, TMEM33 or ENDOD1 did not markedly stabilise RNF26 nor did they alter turnover of immature CD147 (Fig. 5f), a well-characterised Hrd1-dependent ERAD substrate^96^. TMED1 knockdown slowed degradation of both RNF26 and CD147, suggesting it may impact an ER-resident process, such as trafficking, more generally. In contrast to RNF26, its HCIPs appeared stable and were unaffected by loss of their counterparts (Fig. 5f), even though in some cases they were no longer in a complex (Fig. 5e). Of note, VCP/p97 inhibition by NMS-873 only partially restored RNF26 while stabilising CD147 equivalently to MG132 (Fig. 5f, 4a). RNF26 was not among those E3s that enriched VCP/p97 (Fig. 3c), potentially distinguishing its mechanism of turnover from that of a canonical, misfolded ERAD substrate. Neither MG132 nor NMS-873 increased basal levels of RNF26 HCIPs, consistent with relative stability in the ER membrane. These data indicate RNF26 instability is intrinsic, unaffected by its interactions with more stable HCIPs.

### RNF26 interactors regulate STING-dependent innate immune signalling

The immune signalling adaptor STING (<u>ST</u>imulator of <u>IN</u>terferon <u>G</u>enes, also known as MITA/TMEM173) is a polytopic ER-resident protein activated by the cyclic dinucleotide cGAMP, a product of cyclic GMP-AMP-synthase (cGAS) generated in response to the presence of cytosolic double-stranded DNA^97^ (Fig. 6a). Activation of dimerised STING by cGAMP triggers higher order oligomerisation in the ER^98^, subsequent efflux into ERGIC-derived vesicles^99,100^, and type I interferon (IFN) signalling through recruitment of TBK1 and IRF3, leading to its eventual turnover by p62/SQSTM-dependent autophagy^101^. In the ER, STING is a target for ubiquitination by multiple E3s including RNF26^86,102,103^, which are reported to modulate its IFN signalling capability through UPS-dependent degradation. STING and RNF26 reportedly interact through their TMDs and the absence of RNF26 (or its RING domain) counterintuitively increases STING turnover and attenuates IFN signalling^86^. As anticipated, endogenous STING co-precipitated with FH-RNF26 in an interaction reduced following activation by directly adding cGAMP (Fig. 6a), consistent with departure of STING from the ER that reduces its proximity to RNF26. MG132 stabilised RNF26 and consequently restored co-precipitation of STING even in the presence of cGAMP, although concurrent impairment to UPS-dependent degradation of the ER-resident population of STING might also be a contributing factor.

**Fig. 6.**
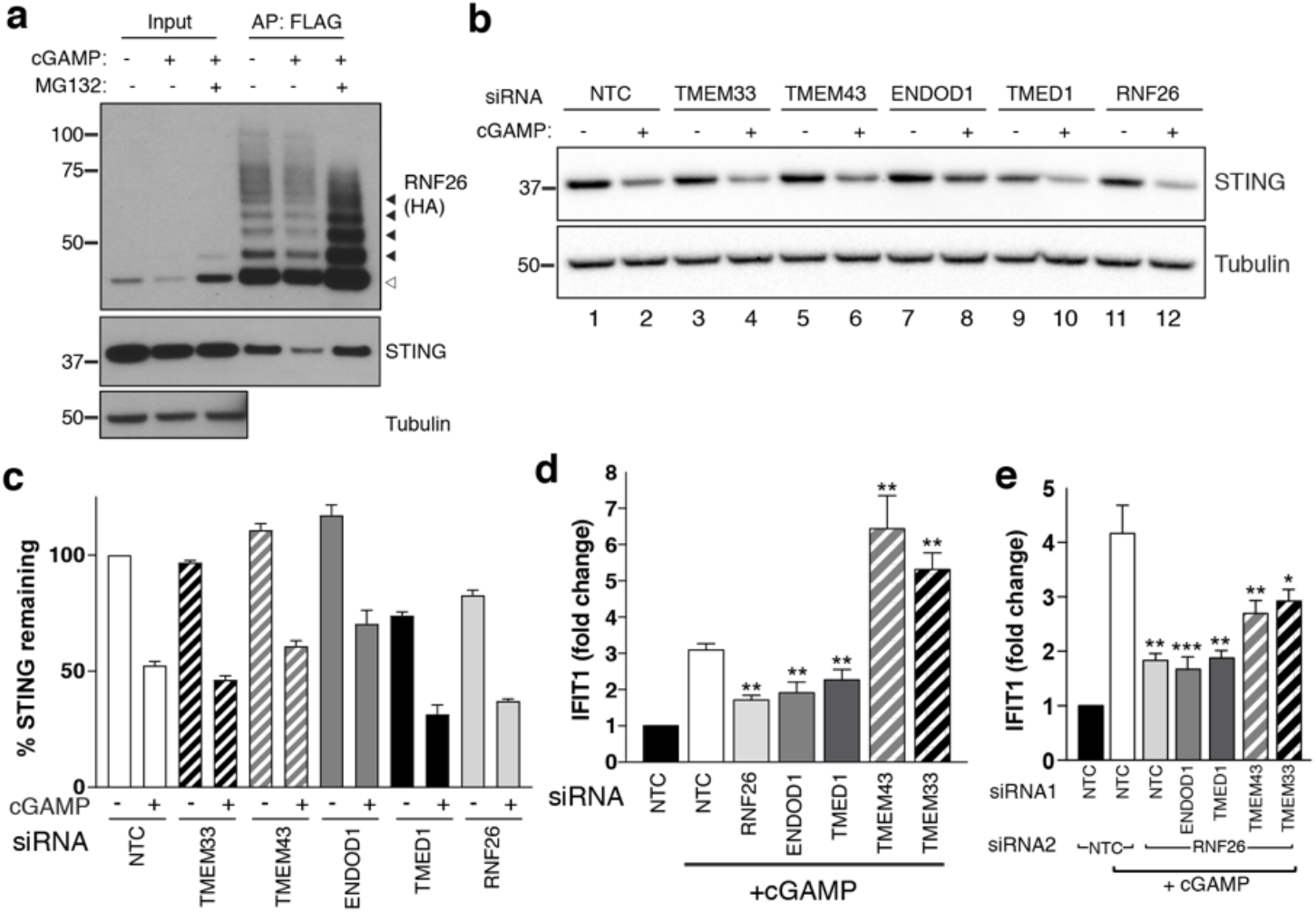
RNF26 and its interactors modulate STING-dependent innate immune signalling. (**a**) WT FH-RNF26-expressing cells treated with cGAMP (6 h, 10 μg/ml) ± MG132 (10 μM, 2 h), with FH-RNF26 isolated from 1% LMNG lysates by anti-FLAG IP, separated by SDS-PAGE and the resulting western blots probed for both RNF26 (HA) and STING. Black arrowheads indicate ubiquitinated forms of RNF26. (**b**) Representative western blot of cGAMP-treated Flp-In^™^293 cells transfected with siRNA targeting RNF26 and HCIPs (TMEM33, TMEM43, ENDOD1, TMED1) and probed for STING and tubulin. (**c**) Quantification of 3 biological replicates for (b) with mean and S.E.M. shown (n=3). (**d**) qRT-PCR for *IFIT1* and *GAPDH* from cGAMP-treated Flp-In^™^293 cells (5 μg/ml, 6 h) transfected with siRNAs targeting RNF26, ENDOD1, TMED1, TMEM43 and TMEM33, along with a non-targeted control (NTC). Normalised *IFIT1* levels in cGAMP-treated cells are shown relative to their untreated counterpart for each siRNA. Mean and S.E.M are shown for at least 4 biological replicates. (**e**) Same as (**d**) but including siRNA targeting RNF26 along with HCIPs. Mean and S.E.M are shown for four biological replicates. For all statistical analysis, \**P* < 0.05, \*\**P* < 0.01, \*\*\**P* < 0.001. Details of statistical analysis are in Extended Data Table 12.

We next considered whether RNF26 HCIPs influenced STING and observed that like RNF26 deficiency, TMED1 siRNAs reduced STING protein levels by ~30% whereas those targeting TMEM43, ENDOD1 or TMEM33 had little to no impact (Fig. 6b,c). Directly introducing cGAMP activates STING and promotes its trafficking and subsequent degradation, reflected by reduced detection of STING protein (Fig. 6b,c). Reductions in STING abundance were proportional in all knockdowns (~50%), indicating that the influence of RNF26 and TMED1 did not depend on whether or not STING could be activated.

To ascertain whether RNF26 HCIPs also influence STING-mediated signalling in response to cGAMP, we monitored transcription of the interferon-stimulated gene (ISG) *IFIT1* (interferon-induced protein with tetratricopeptide repeats 1) after silencing HCIPs alone and together with RNF26. STING activation by cGAMP increased *IFIT1* transcription ~3-4 fold in WT cells (Fig. 6d,e) and did so in a dose-dependent manner (Extended Data Fig. 6a). RNF26 knockdown (Fig. 6d, Extended Data Fig. 6b) attenuated the increase in *IFIT1,* consistent with previous findings^86^. Expression of FH-RNF26_Y432A_ also inhibited *IFIT1* upregulation (Extended Data Fig. 6c), demonstrating that the *IFIT1* response requires ubiquitination by RNF26 and the Y432A mutant functions as a dominant negative. Knockdown of either TMED1 or ENDOD1 also dampened the cGAMP-induced *IFIT1* response (Fig. 6d), but lower STING levels only coincided with TMED1 loss and not ENDOD1 (Fig. 6b,c), suggesting these two HCIPs impact STING and signalling through it, by different mechanisms. In marked contrast, depleting cells of either TMEM43 or TMEM33 enhanced *IFIT1* responses to cGAMP, resulting in 5-7-fold increases that were nearly twice the magnitude elicited from CTRL cells (Fig. 6d). These results were recapitulated with an independent set of siRNAs (Extended data Fig.6d). Co-depleting RNF26 along with TMED1 or ENDOD1 did not attenuate *IFIT1* responses further, and co-depleting with either TMEM43 or TMEM33 reduced the enhanced *IFIT1* responses by ~50% (Fig. 6e), indicating that the impact of these HCIPs on STING activation was at least in part through RNF26. These data describe four novel modulators of STING-dependent immune signalling and define the RNF26 complex as an immunoregulatory unit.

## Discussion

The ER accommodates a range of functional modalities; many of which fall under the regulatory remit of ubiquitination by resident E3-containing complexes. Comparative proteomic strategies have generated extensive and informative interaction networks of ubiquitination machinery^51,104^. We have applied those principals to expand the functional ERAD modules described for Hrd1 and gp78/AMFR^24^ by mapping over 450 interactions to form the landscapes for 21 ER-E3s. The ER membrane is increasingly appreciated as a diverse site of regulation for metabolic and homeostatic processes, which is reflected in the diversity of interactions made by resident E3s.

### HCIPs represent cofactors and substrates of ER-E3s

ER-E3s assemble multi-subunit complexes from protein-protein interactions made through lipid bilayer, lumenal, and cytosolic contacts, making these units highly adaptable at conjugating Ub proximal to the membrane. Because ER-E3s appear structurally and topologically diverse, the minimal redundancy their HCIP networks exhibit (Fig. 2a) is consistent with assembling complexes from dedicated cofactors that would favour selective cellular responsibilities. More than 70% of HCIPs identified contain predicted TMDs or signatures associated with ER residency, among which will be both E3 cofactors and substrates. E3 cofactors are exemplified by Hrd1 interactors such as SEL1L and FAM8A1, but also by those validated for RNF26 (Fig. 5) and proposed for RNF185 (Extended Data Fig. 3b). Substrates are typified by the Wnt receptor Evi/WLS/GPR177, which after initially being identified early in this study was established as novel target for regulated ERAD by CGGRF1^105^. Other promising candidate substrates might include DHCR7, the terminal enzyme of cholesterol biosynthesis found with Hrd1 and a validated ERAD substrate^106,107^ and ACSL4, an RNF185 HCIP catalysing long chain polyunsaturated CoA synthesis^108^ which global turnover studies indicate has a short half-life^109^. SLC39A14/ZIP14, a stress-regulated Zn^2+^ and Mn^2+^ transporter at the plasma membrane, was identified as a Hrd1 HCIP. As it was detected under ER-stress free conditions, Hrd1 may be responsible for constitutive degradation of ZIP14 to maintain metal homeostasis. Like these, additional E3-substrate relationships are likely to be present within the dataset but will require detailed kinetic analyses to confirm them. Identifying and differentiating cognate cofactors from substrates of E3s remains a significant challenge in ubiquitin biology. Proteomics-based studies such as this, cannot readily distinguish E3 substrates from cofactors participating in substrate recruitment, complex assembly, localisation or ubiquitin conjugating activity. As the example of RNF26 HCIPs illustrates, biochemical, genetic and functional validation remains essential to accurately define complexes within each individual network. Although not providing comprehensive insight into all ER-E3 complexes that form, our analysis establishes a starting point from which to systematically elucidate their molecular organisation and functional responsibilities.

### Ubiquitin linkages and machinery associated with ER-E3s

Ubiquitination at the ER is often envisaged in terms of conventional ERAD, where associated ubiquitin linkages (e.g. K48, K11) added to substrates by one (or more) ER-E3 facilitates recognition by ubiquitin-binding proteins associated with VCP/p97^110^ and proteasomes^111^. But our findings highlight the potential of some resident E3s to function differently. For example, RNF26 reportedly modifies STING with K11 chains^86^ but our data demonstrate its own turnover from the ER is rapid and is associated with K33-, K63- and/or K29-linked ubiquitin chains not conventionally associated with degradation or targeting to proteasomes. K29-linkages can be part of heterotypic branched and mixed Ub chains^112^, which can hallmark some ERAD substrates^113^. What role linkage heterogeneity plays in RNF26 turnover is not yet clear but might indicate linkages other than K48- and K11- also possess the ability to target proteins from the ER to proteasomes^111^.

E2s are determinants of poly-ubiquitin linkages while associated DUBs trim and prune them with linkage-specificity, yet most pairwise relationships with E3s remain undefined. As there are ~40 E2s and nearly ~100 DUBs in the human proteome, we would expect one (or more)to partner with each E3 to expand its substrate range. Envisioning ER-E3 prey profiles that enriched UPS components, we instead found that E2s and DUBs were not prominent among HCIPs. While key interactions for ERAD were confirmed (UBE2J1-Hrd1^24,48^ and UBE2G2-gp78^114^), the often transient, low affinity E3-E2 and E3-DUB interactions appear not to have been widely preserved. Yeast-2-hybrid screens used to define human E2-E3 pairs have reported ER-E3s (e.g. RNF26, BFAR, RNF5) as interactors of multiple E2s from prey libraries^115,116^, but these have yet to be validated *in vivo* and were not identified as HCIPs. Similarly, functional roles in ERAD were identified for USP13, Atx3, VCPIP and USP19, but ER-E3s were not identified among the DUB interactome^51^. Detailed mapping of the ubiquitin linkage landscape attributable to individual ER-E3s and the UPS machinery required, remains an outstanding question for future studies.

### The specialised role of Hrd1 among ER-E3s

ERAD is responsible for misfolded protein disposal during ER stress to restore organelle homeostasis. Induction of any ER-E3s by pro-survival UPR branches might have signalled complementary contributions to ERAD and ER stress resolution provided by the Hrd1 complex. Although ER stressors upregulated some ER-E3s (e.g. CGRRF1, RNF170, RNF5; Fig. 3b,^57^), their upregulation was modest relative to that observed for Hrd1. This is consistent with the observation that Hrd1 was the only E3 upregulated by direct activation of either the XBP1 or ATF6 transcription factors^117^. Moreover, there were only 24 HCIPs among known UPR target genes^58^ with at least a quarter belonging to the Hrd1 complex (Extended Data Table 7,^24^). Thus, evolutionary expansion of E3s in the mammalian ER has not extensively supplemented adaptive stress resolving capabilities functionally redundant with Hrd1, consistent with ubiquitination by the Hrd1 complex being essential for survival under proteotoxic ER stress conditions^60,118^. With misfolding of secreted rather than membrane proteins being the principal instigators of ER stress, an exclusivity of access to lumenal substrates via SEL1L offers an explanation why Hrd1 appears indispensable. Instead, other ER-E3s already known to oversee specific metabolic or regulatory functions consistent with higher order organismal functions, appear more likely to have evolved more selective client ranges of integral membrane proteins. ER-E3s may only hold strategic importance in selected cells or tissues, where a particular substrate/s are physiologically relevant. The enrichment of VCP/p97 only by only some ER-E3s (e.g. Hrd1, gp78, RNF185) supports a model where ERAD-like degradation is only one of the processes E3s oversee at this membrane interface.

### Protein-conducting channels within ER-E3 complexes

An essential facet of ERAD is delivery of misfolded ER proteins to cytosolic proteasomes. Recent Cryo-EM structures containing recombinant yeast Hrd1p compellingly show the presence of an aqueous cavity formed by its TMDs that could provide a retrotranslocation conduit for misfolded polypeptide transport across the ER membrane^15,119^. With functionally demonstrated roles in ERAD of membrane proteins, other ER-E3s might be expected to form structurally analogous protein-conducting channels to dislodge substrate TMDs. Some ER-E3s (e.g. MARCH6, Trc8/RNF139) encode a sufficient number of TMDs from which such a channel could reasonably be formed from monomers, while others (e.g. RNF185) would require higher-order oligomers or large multi-subunit, TMD-containing HCIP/cofactor assemblies. If analogous cavities are formed, the absence of HCIPs common to ERAD-related E3s might indicate the evolution of specialised strategies to selectively remove TMD-containing substrates. So far only Hrd1 has been implicated in “retrotranslocation” of misfolded lumenal proteins, whereas multiple ER-E3s (e.g. MARCH6, Trc8/RNF139, TMEM129, gp78/AMFR) as well as Hrd1, appear sufficient to “dislocate” TMDs of integral membrane substrates^33,37,46,47,120^. Perhaps only the Hrd1 protein-conducting channel is optimised to accept lumenal substrates, thus providing a key function for ER stress resolution. The ER’s lipid bilayer presents a formidable barrier for downstream degradation machinery. With lateral access, membrane-bound substrates seem poised to access an ER-E3 conduit differently or engage proteases to cleave TMDs and release fragments. Perhaps this is why coordinated involvement of intramembrane proteases such as SPP or the rhomboid RHBDL4 in ERAD has always been conceptually appealing^67,121^. In all kingdoms of life, membrane-spanning protein complexes are capable of translocating polypeptides represent specialised, macromolecular apparatuses^122^. If and how ER-E3s mediate this process is a rich area for future exploration.

### RNF26 is an intrinsically unstable ER-E3

We found that RNF26 auto-ubiquitination coincided with its rapid turnover from cells (Fig. 4c). Sequence similarity between the RNF26 and RNF4 C-termini, notably in a conserved penultimate Tyr residue required for Ub transfer (Fig. 4b), explains its recalcitrance to detection at the protein level. RNF4 homo-dimerisation allows activated E2 to bind one monomer while engaging the other to activate Ub-transfer to substrate. The Ub Ile44 hydrophobic patch binds Tyr189 at the RNF4 dimer interface, permitting the E2-Ub oxyester to be efficiently hydrolysed^88^. Evidence of dimerisation along with the observation that the Y432A mutant phenocopies ubiquitination defects like those observed when directly disrupting the RING domain or E2 binding (e.g. C401S, I382R^41,86^), suggests that RNF26 embraces a Ub conjugating mechanism like that of RNF4 and related E3s (e.g. IAPs).

### STING regulation by RNF26

Chronic activation of STING is linked to autoimmune and auto-inflammatory disorders^123^, denoting an imperative for tight regulation and fine control over this signalling cascade. Ubiquitination offers complex spatiotemporal, post-translational control of STING abundance, activation and consequently, signalling. Multiple E3s, including RNF26, reportedly ubiquitinate STING while in the ER and after it is trafficked into endolysosomal vesicles following cGAMP-induced oligomerisation^98^. RNF26 knockdown lowered STING levels and dampened *IFIT1* upregulation (Fig. 6b,c,d). Expression of FH-RNF26_Y432A_ exerted the same effect on *IFIT1* levels (Extended Data Fig. 6c), placing RNF26-mediated ubiquitination as a potent modulator of IFN signalling^86^. One model posits that RNF26 competes with RNF5 to extend K11-rather than K48-Ub chains on Lys150 of STING^86^, where dynamic balancing of ER-E3 ubiquitination reactions governs STING proclivity for degradation, based on affinities of Ub-binding proteasome subunits for different linkages. Lys150 lies in the connector helix loop just after the final TMD (TM4,^98^) and so would be accessible by different ER-E3s. Domain mapping demonstrated RNF26 and STING interact through TMDs^86^, but further investigation is required to clarify how RNF26 might ubiquitinate STING and not itself, perhaps by recruiting a substrate-selective cofactor (e.g. an HCIP) or by engaging different E2s. We found RNF26 capable of modifying itself with K63-, K33-, and/or K29-Ub chains and not K48- and K11- as shown previously for STING^86^, which distinguishes the ubiquitination reactions occurring in cis- and trans- by RNF26 and raises the potential involvement of different E2s.

### An RNF26 complex scales the IFN response through STING

We identified four RNF26 interactors capable of scaling the IFN response through STING. These modulators were not found previously by RNF26 proteomics, which only used the RING-containing, cytoplasmic C-terminus as bait^41^. Together with RNF26, TMEM43, TMEM33, ENDOD1, and TMED1 form a membrane-bound complex that appears capable of influencing STING activation, most likely through its membrane-spanning TMDs. Although these RNF26 HCIPs do not appear structurally similar or functionally orthologous, they share the ability to scale IFN signalling through cGAS-STING pathway in an RNF26-dependent manner (Fig. 6e), suggesting they act collectively through one or more complexes. How each RNF26 HCIP accomplishes this is not yet clear, as their individual functions are not yet fully appreciated.

Loss of TMEM43 or TMEM33 increases the IFIT1 response (Fig. 6), consistent with roles as negative regulators of STING activation. As TMEM43 was required for RNF26 interaction with TMEM33 (Fig. 4e), enhanced signalling through STING could be explained by the loss of TMEM33 from the RNF26 complex. TMEM33 is the human ortholog of *S. pombe* Tts1, an ER-shaping protein that helps sustain high-curvature ER membranes^124^ and is involved in nuclear envelope remodelling during mitosis^125^. Functional conservation in metazoans could indicate that local ER membrane curvature is an important determinant of STING activation. Alternatively, its role in organising the peripheral ER and as a binding protein of reticulons could mean that TMEM33 influences localisation of RNF26 and STING throughout the ER/INM network, which then determines signalling capability and regulation. TMEM33 has been linked recently to regulation of intracellular calcium homeostasis through an interaction with polycystin 2 (PC2), suggesting its influence extends beyond its contribution to a complex that modulates STING or RNF26. TMEM43/LUMA localises in both the INM and the ER and interacts with proteins such as Lamin A/B and Emerin^92^. Mutations in TMEM43 are genetically linked to the heritable cardiomyopathy autosomal dominant arrhythmogenic right ventricular cardiomyopathy/dysplasia (ARVC/D, S358L)^126^ and the autosomal recessive myopathy Emery-Dreifuss Muscular Dystrophy (EDMD, Q85K, I91V,^127^). Whether these rare conditions are in some way attributable to the suppression of STING activation by TMEM43 mutants is not known. TMEM43 has been linked previously to immune signalling and NF-kB through an interaction with CARMA3/CARD10 and EGFR^128^, but not previously to STING.

Silencing TMED1 or ENDOD1 phenocopied the reduction of *IFIT1* levels upon cGAMP treatment that is observed with loss of RNF26, consistent with both HCIPs functioning to either permit or enhance STING activation. TMED1/tp24 contains a GOLD domain and is part of the p24 family of trafficking proteins (TMED1-10). It forms monomers and dimers instead of hetero-oligomers with other p24 family members for function^94^. TMED1 may influence vesicle trafficking from the ER as a cargo receptor or it may be linked to vesicle coat formation, potentially placing it as a gatekeeper for budding ER vesicles containing proteins such as STING. Notably Kelch-like 12 (KLHL12), a CUL3 adaptor and protein linked to ER-Golgi trafficking through regulation of COPII vesicle size^129^, was an HCIP enriched by RNF26_Y432A_. ENDOD1 is predicted to contain a non-specific endonuclease domain shown to have nuclease activity *in vitro* using an orthologous domain from *Paralichthys olivaceus* (Japanese flounder)^130^. Importantly, this study identified ENDOD1 among the genes that are important for innate immunity in fish, suggesting a role that is evolutionarily conserved. It is not yet clear from our study whether ENDOD1 and STING interact directly or what may be the implications of having endonuclease activity near a hub for immune signalling.

Our data reveal that RNF26 nucleates an immuno-regulatory complex, which was discovered through constructing the interaction landscape of the ER-resident E3s. Information within this landscape provides a resource to uncover E3 functions within various cellular processes. Understanding the mechanisms modulating abundance and activity of its membrane-embedded proteins will help to determine if they represent tractable targets that may be leveraged for potential therapeutic benefit.

## Methods & Materials

### Plasmids and Transfections

All cDNAs encoding individual ER-E3s (Extended Data Table 9) were amplified and appended with restriction site-containing linkers by PCR, and subsequently subcloned into a pcDNA^™^/5/FRT/TO vector (Invitrogen) containing a FLAG-HA (FH) tag in frame (N- or C-terminal) by restriction digest and ligation. Sequences for HCIPs (Extended Data Table 10) were obtained and processed similarly but subcloned instead into pcDNA3.1(-) vectors containing either an N- or C-terminal S-tag in frame as described previously^27^. All plasmids were transfected into recipient cell lines using Lipofectamine^™^2000 (Thermo Fischer Scientific) according to manufacturer’s guidelines.

### Cell culture and generation of stable cell lines

Flp-In^™^ T-REx^™^ human embryonic kidney 293 cells (Thermo Fischer Scientific, referred to as Flp-In^™^293) were used to generate stable cell lines individually expressing selected E3s. Briefly, each FH-E3-pcDNA5/FRT/TO construct was co-transfected with the Flp recombinase vector pOG44 (3:1 ratio) as described above. Cell pools stably recombining and expressing E3s were selected by resistance to Hygromycin B (100 μg/ml, InvivoGen). All Flp-In^™^293 cell lines were cultured in DMEM (Lonza, BE12-604F) supplemented with 10% (v/v) fetal bovine serum (FBS) and glutamine (2 mM). All cells were grown at 37°C and 5% CO_2_.

### Antibodies and compounds

The following primary antibodies were used for detection by Western blot: anti-Hrd1 (Bethyl, #A302-946A), anti-SEL1L (Santa Cruz Biotechnology, #sc-48081), anti-UBE2J1 (Abcam, #ab39104), anti-OS-9 (kind gift from R. Kopito, Stanford), anti-Herp (Abcam, #ab150424,) anti-Derlin1 (kind gift from Y.Ye, NIH), anti-tubulin (Sigma, #T6074), anti-FLAG (Sigma, #F3165 and #F7425), anti-HA (Sigma, #H9658; Cell Signaling Technologies, #3724) anti-ubiquitin (Cell Signaling Technologies, #3933 and #P4D1), anti-S-tag (Thermo Scientific, #MA1-981), anti-TMEM43 (Abcam, #ab184164), anti-TMED1 (Abcam, #ab224411), anti-ENDOD1 (Abcam, ab121293), anti-AUP1 (Atlas Antibodies, #HPA007674), anti-UBXD8 (Proteintech, #16251-1-AP), anti-STING (Cell Signaling Technologies, #D2P2F), anti-CD147 (Santa Cruz Biotechnology, #sc-25273). Anti-FAM8A1 has been reported previously^27^. The secondary antibodies used for western blot and IF include: goat anti-rabbit HRP (1:10,000, BioRad), goat anti-mouse HRP (1:10,000, Santa Cruz Biotechnology), donkey anti-goat HRP (1:10,000, Santa Cruz Biotechnology), goat anti-rabbit-Alexa 488 (1:400, Life Technologies), goat anti-mouse-Alexa 568 (1:400, Life Technologies). The following compounds were used in this study; MG132 (10 μM, Merck Millipore), Tunicamycin (500 ng/ml, Sigma), NMS-873 (10 μM, Sigma), cycloheximide (100 μg/ml, Abcam), N-ethylmaleamide (NEM, Acros Organics), DAPI (Sigma), doxycycline (DOX, Sigma), dithiolthreitol (DTT, Sigma), iodoacetamide (IAA, Roche) and SubAB^56^.

### siRNA transfections

Flp-In^™^293 cell lines seeded in 12-well plates were transfected with individual siRNAs (50 nM, Sigma, Extended Data Table 11) using Lipofectamine RNAiMax (Thermo Fischer Scientific) at a ratio of 1:4 according to the manufacturer’s instructions. Cells were expanded 24 h post-transfection and harvested following another incubation of 24 h.

### Mass Spectrometry and Proteomic analysis

Each FH-E3-expressing Flp-In^™^293 cell line was seeded in 15 cm plates and treated with DOX (1-1000ng/ml, 18 h). Cells were harvested at ~80% confluence, washed and subsequently resuspended in solubilisation lysis buffer (SLB: 150 mM NaCl, 50 mM Tris-HCl pH7.4, 5 mM EDTA) containing 1% Lauryl Maltose Neopentyl Glycol (LMNG, Anatrace) and supplemented with cOmplete^™^ protease inhibitor cocktail (Roche). Lysates were clarified by centrifugation (20,000 x g, 30 min.) and pre-cleared using CL-4B Sepharose beads (50μL of 50:50 slurry, Pharmacia/GE). The resulting clarified whole cell lysate (WCL, 10 mg) was used as source material for immunoprecipitations with anti-FLAG agarose (M2, Sigma, A2220) for 2 h. Immunoprecipitated complexes were washed twice with SLB (without detergent), twice with TBS, and eluted by 2 x SDS + 10% β-Mercaptoethanol. Eluates were reduced by DTT, alkylated by IAA, and subject to double chloroform-methanol precipitation. Precipitated proteins were subject to tryptic digest prior to purification using C18 Sep-Pak cartridges (Waters). Purified peptides were analysed by LC-MS/MS with a tandem mass spectrometer (Q Exactive^™^ HF, Thermo Fischer Scientific) with an EASY-Spray^™^ C18 LC Column (2 μm, 100Å, 75 μm x 50 cm, Thermo Fischer Scientific) over a 63 min 2-35% acetonitrile gradient in 5% DMSO (v/v)/0.1% formic acid (v/v). The data were acquired with a resolution of 70,000 full-width half maximum at mass/charge 400 with lock mass enabled (445.120025 m/z), Top 15 precursor ion selection, Dynamic Exclusion of 27 s, and fragmentation performed in Higher-energy C-trap dissociation (HCD) mode with Normalized Collision Energy of 28. Samples were analysed twice to generate technical duplicates. Chromatogram alignment and peptide intensity were determined by Progenesis-QI (nonlinear Dynamics). Peptides were identified and matched using the SwissProt database. Each bait sample was assigned with more than 2,000 protein IDs. Assignment of p-values to identified proteins was accomplished by adapting the comparative BSCG method described previously^50^. High-confidence interacting proteins (HCIPs) were defined by an ability to meet four criteria: 1) p-value <0.05, 2) positive foldchange, 3) identified by >1 peptide, and 4) not classified as ‘common’ contaminants (Extended Data Table 3). SINQ analysis^52^ was also carried out to identify interactors unique to one E3 only, which would not be assigned a p-value with the comparative analytical method. Raw data have been deposited in the PRIDE database (submission ongoing).

### Immunoprecipitation, SDS-PAGE and western blotting

Cells rinsed in phosphate-buffered saline (PBS) were mechanically lifted, harvested, and lysed in SLB + 1% LMNG or Triton X-100 (TX-100, Fischer Scientific), as described above. Lysates were clarified by centrifugation (17,000 x g, 30 min.) and pre-cleared using CL-4B Sepharose beads (50μL of 50:50 slurry, Pharmacia/GE), with subsequent affinity and immunopurifications carried out using the resulting lysates. Beads were washed thrice in SLB and subsequently resuspended in 2 x Laemmli buffer + 20 mM DTT after the final wash, separated by SDS-PAGE and transferred to PVDF membrane for western blotting. Western blots were performed by incubating membranes in PBST blocking buffer (PBS + 1% Tween-20 supplemented with 5% non-fat dry milk), with subsequent primary and secondary antibody incubations in PBST + 5% non-fat dry milk. Secondary antibodies conjugated with horseradish peroxidase (HRP) were used to detect proteins bound to primary antibodies for enhanced chemiluminescence (ECL) with images captured either on X-ray film (FujiFilm, SuperRX) or by CCD camera (Chemidoc, BioRad) for quantification.

### Quantitative transcript analysis by NanoString

RNA from Flp-In^™^293 cells was extracted using the RNeasy kit (Qiagen) in accordance with the manufacturer’s instructions that included the genomic DNA digestion with DNaseI (Qiagen). Isolated RNA from each sample (150 ng) was hybridized to a Reporter CodeSet and Capture ProbeSet (10μL each) for a selected set of genes (Extended Data Table 6) by incubating in hybridization buffer (65°C, 18 h) and loaded in the nCounter^®^ PrepStation according to manufacturer’s instructions. Hybridized probe/target complexes were immobilized on the nCounter^®^ Cartridge and imaged in the nCounter^®^ MAX Digital Analyzer (high-resolution setting). Data were processed and analysed according to the manufacturer’s guidelines (NanoString Technologies Inc.). All experiments were performed in biological triplicate (n=3).

### Radiolabelling and pulse-chase

Radiolabel pulse-chase assays of Flp-In^™^293 cells stably expressing either FH-RNF26_WT_ or FH-RNF26_Y432A_ were carried out as previously described^22,27^. Briefly, following DOX treatment (18 h) cells were starved in DMEM (Lonza) lacking methionine (Met) and cysteine (Cys) + 10% dialysed FBS for 10 min, metabolically labelled by supplementing starvation medium with^35^S-Met/Cys (EXPRE^35^S^35^S Protein Labelling Mix (PerkinElmer), 80 μCi/6 cm plate) for 10 min, rinsed thrice in PBS, and chased for indicated time points in DMEM supplemented with Met and Cys (50 mM each). Cells were lysed in SLB containing 1% TX-100, and the detergent-soluble, post-nuclear lysates pre-cleared using CL-4B Sepharose beads followed by immunoprecipitation with anti-HA-antibody (12CA5) and Protein G agarose (Roche). Bead-bound radiolabelled substrates were resuspended in 2 x Laemmli buffer (+20 mM DTT), separated by SDS-PAGE and imaged using a phosphoimager (BioRad).

### cGAMP transfection for *IFIT1* qRT-PCR

Flp-In^™^293 cells seeded in 24-wells were stimulated by transfecting 5 μg/ml cGAMP using Lipofectamine 2000 at a ratio of 1.25:1. Cells were harvested 6 h post-transfection and the extracted RNA (RNeasy, Qiagen) reverse transcribed to produce cDNA (QuantiTect, Qiagen) according to manufacturer’s instructions. Taqman probes targeting human *GAPDH* (Hs02758991_g1) and *IFIT1* (Hs03027069_s1) were purchased from Life Technologies. qRT-PCR data were collected on a StepOnePlus^™^ Thermal Cycler (Thermo Fischer Scientific) and analysed by the ΔΔC_t_ method, normalising *IFIT1* levels to GAPDH. Averages and S.E.M. were determined from triplicate assays from at least three independent experiments (n=3).

### Immuno- and affinity purification of ubiquitinated proteins

FH-RNF26_WT_ and FH-RNF26_Y432A_ -expressing Flp-In^™^293 cells were induced with DOX (18 h) and where indicated, samples were additionally treated with 10 μM MG132 for 2 h. Cell pellets were lysed in TUBE lysis buffer (20 mM sodium phosphate pH 7.5, 1% NP-40 (v/v), 2 mM EDTA, supplemented with cOmplete protease inhibitor cocktail (Roche), PhosSTOP (Roche), NEM (50 mM), and DTT (1 mM)). Lysates were centrifuged (as above) and the detergent-soluble fraction subsequently incubated overnight with 15 μL magnetic GST resin (Thermo Fischer Scientific) conjugated to 50 μg 1x UBA-His_6_ binder. Bound resin was washed thrice with TUBE lysis buffer and each sample split equally to accommodate control/untreated or USP21 deubiquitinase treatment.

### Deubiquitination assays

Bead-bound material from UBA-His_6_ binder pulldowns (above) was resuspended in deubiquitinating-buffer (50 mM HEPES pH 7.5, 100 mM NaCl, 2 mM DTT, 1 mM MnCl_2_, 0.01 % Brij-35) without or with 0.5 μM USP21 (Ubiquigent). Samples were incubated (1 h, 30°C) in a thermoshaker (VWR, 750 rpm) and subsequently denatured by incubation (65°C, 20 min) with 2 x Laemmli buffer + DTT (20 mM) followed by separation on SDS-PAGE. For UbiCREST analysis^131^, FH-RNF26_WT_ from stably expressing Flp-In^™^293 cells (4mg) was immunoprecipitated by anti-FLAG agarose, washed, divided and individually incubated with the panel of recombinant dUbs, according to the manufacturer’s instructions (Boston Biochem).

### I*n vitro* ubiquitination assay

FH-RNF26_WT_ and FH-RNF26_Y432A_ -expressing Flp-In^™^293 cells were treated and lysed in TUBE lysis buffer as described above. Resulting supernatants were incubated with anti-FLAG M2 magnetic beads (Sigma, 3 h at 4°C). Beads were washed thrice with TUBE lysis buffer, dividing samples in half prior to the last wash and then washed once with *in vitro* ubiquitination (IVU) buffer (50 mM Tris-HCl, pH 7.5, 2.5 mM MgCl_2_). Assays were carried out by resuspending beads in IVU buffer supplemented with; ubiquitin (10μM, Boston Biochem), E1 enzyme (150 nM, Enzo), UbcH5a (1 μM, Enzo), and DTT (0.5 mM), with or without ATP (4 mM, pH 8.0, Sigma) and incubated in a thermoshaker (15 min, 30°C, 750 rpm). Samples were denatured with 2 x Laemmli buffer + 20 mM DTT (65°C, 20 min).

### Velocity sedimentation

Velocity sedimentation was carried out as previously described^27^. Briefly, DOX-induced Flp-In^™^293 cells (1μg, 18 hrs) were mechanically harvested and lysed in SLB containing 1% LMNG (as described above). Post-nuclear, pre-cleared WCLs (1 mg total) were layered onto either a continuous sucrose gradient (10-40% or 5-30%) prepared using a Gradient Master 108^™^ (BioComp). Sucrose was dissolved in a physiological salt solution (150 mM NaCl, 50 mM Tris-HCl pH 7.4, 5 mM EDTA, 1 mM PMSF) + 1% LMNG. Samples were centrifuged in an SW.41 rotor (OptimaTM L-100 XP, Beckman Coulter, Brea, CA) at 39,000 rpm for 16 h at 4°C. Thirteen fractions (940 μL each) were collected manually and proteins precipitated by addition of 190 μL 50% (v/v) trichloroacetic acid (TCA). Following acetone washes, precipitated proteins were resuspended in 2 x Laemmli buffer + 20 mM DTT and separated by SDS-PAGE. If necessary, samples were neutralised with 1 M Tris-HCl (pH 9). All samples were heated (10 min, 56°C) and separated by SDS-PAGE. Gel filtration standards (Gel Filtration Markers Kit, MWGF1000, Sigma Aldrich) were separated on similar gradients to estimate protein complex size and included: alcohol dehydrogenase (150 kDa), β-amylase (200 kDa), apoferritin (443 kDa) and thyroglobulin (663 kDa). Standards were processed as above and detected by Coomassie staining.

### Immunofluorescence and microscopy

For detection of FH-E3s, Flp-In^™^293 cells were seeded onto 13 mm poly-L-lysine coated cover slips and induced with DOX (18 h). Cells were fixed with 4% paraformaldehyde (PFA, 20 min at room temperature (RT)), permeabilised with PBS containing 0.2% TX-100 for 5 min at RT and blocked with PBS containing 0.2% PBG (fish skin gelatin) for 30 min. Coverslips were incubated with 1° antibodies diluted in 0.2% PBG (1 h, RT), rinsed twice in PBS and incubated with fluorescent 2° antibodies (0.2% PBG in PBS, 1h, RT). Coverslips were incubated with DAPI (5μg/mL, 10 min, RT) and mounted using ProLong^®^ Gold antifade reagent (Life Technologies). All images were captured on a Zeiss LSM710 confocal microscope and processed in Image J (NIH) and Photoshop (Adobe).

### Statistical analysis

Statistical significance within NanoString data was determined using multiple t-tests (Holm-Sidak method, α = 0.05) that compared fold-change in E3 transcripts from untreated and Tunicamycin-treated (Tm, 500 ng/ml, 8 h) or SubAb5-treated (10ng/ml, 8h) samples. All other data (e.g. qPCR, protein quantification) were analysed using a two-tailed paired t-test comparing non-targeting control siRNA (siNTC) to each siRNA target. All statistical analyses were carried out and plotted using GraphPad Prism (Version 7.0). Detailed statistical information is available in Extended Data Table 12.

### Bioinformatic analysis

Primary amino acid sequences for all E3s and HCIPs were obtained from UniProt (http://www.uniprot.org/), with common motifs annotated using Pfam (http://pfam.xfam.org/)^132^, TMDs predicted by TOPCONS (http://topcons.net/)^133^ and N-linked glycosylation sites predicted by NetNGlyc 1.0 (http://www.cbs.dtu.dk/services/NetNGlyc/). E3 interactions were compared against those previously reported in BioGRID (https://thebiogrid.org/).

## Acknowledgements

We are grateful to Dr. Norbert Volkmar and Dr. Dönem Avci for critical discussions. We also thank Dr. Jan Rehwinkel for technical assistance. E.F. was supported by a fellowship from the Medical Research Council. P.D.C was supported by an EPSRC grant (nr EP/N034295/1) and by the Chinese Academy of Medical Sciences (CAMS) Innovation Fund for Medical Science (CIFMS), China (grant number: 2018-I2M-2-002), awarded to B.M.K. J.C.C. was supported by a grant from the Medical Research Council (MR/L001209/1) and by the Ludwig Institute for Cancer Research.

## Author contributions

E.F. and F.L. designed, performed and analysed experiments; R.F., M-L.T. and B.M.K. performed LC-MS/MS; P.D.C analysed LC-MS/MS raw data; M.G.H. provided reagents and designed cGAMP-STING experiments; A.W.P. and J.C.P provided reagents; K.B. performed *in vitro* ubiquitination experiments; J.C.C. designed, performed and analysed experiments and wrote the manuscript.

---

**Extended Data Fig. 1.**
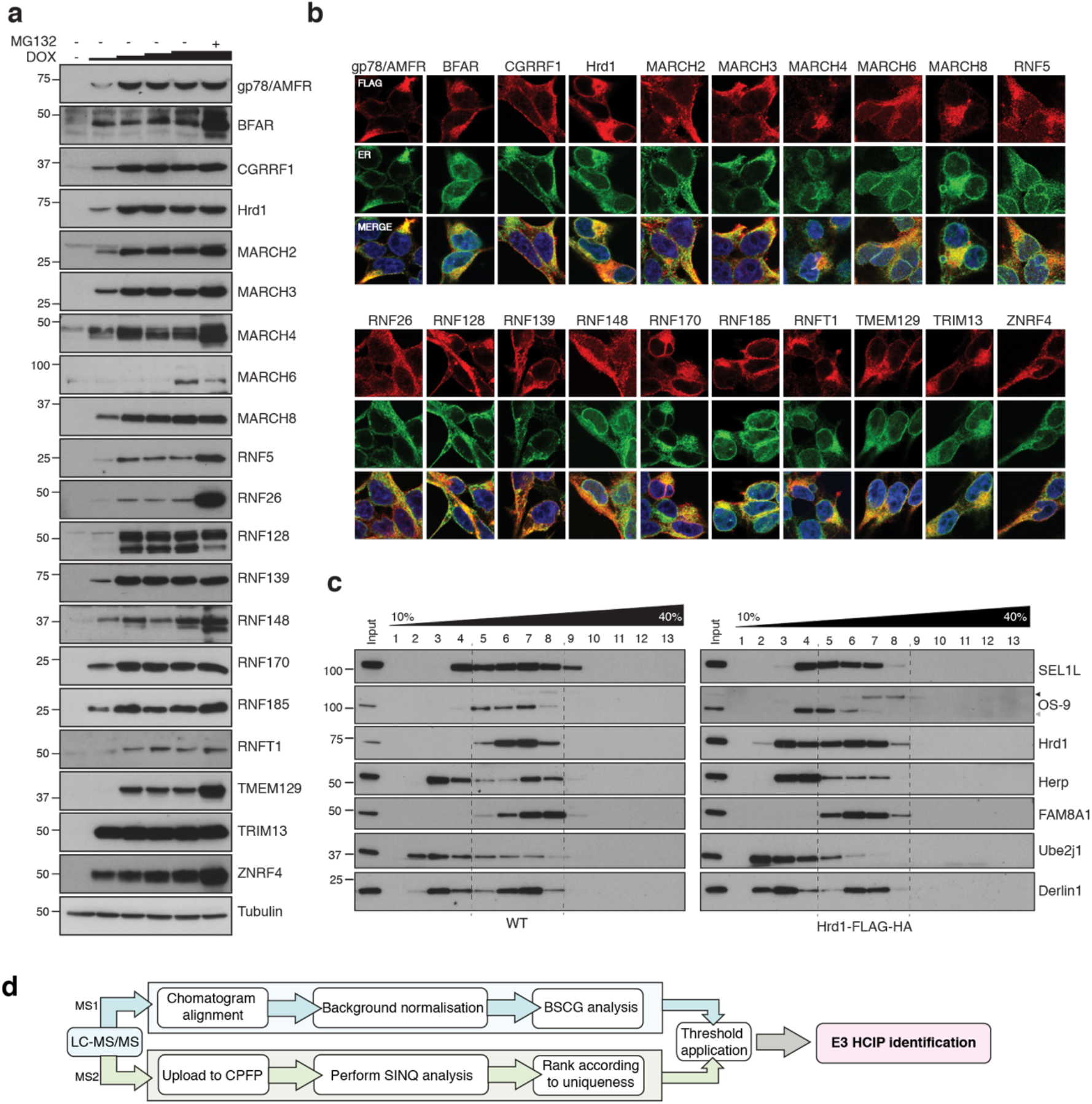
Expression and localisation of FLAG-HA-E3s. (**a**) Doxycycline (DOX) titrations (1, 10, 100, 1000 ng/mL; 18 h) to optimise expression conditions of individual FH-E3s in stable Flp-In^™^293 cell lines. MG132 (10 μM, final 4h) was also included in one sample (DOX, 1000 ng/mL) to evaluate proteasomal degradation. Cells were lysed in RIPA buffer (50 mM Tris-HCl pH7.4, 0.1% SDS, 1% sodium deoxycholate, 1% NP-40, 2 mM EDTA, 150 mM NaCl) supplemented with cOmplete^™^ protease inhibitor cocktail (Roche), lysates separated by SDS-PAGE and western blots probed for each E3 (anti-FLAG). (**b**) Representative immunofluorescence images for each of the generated Flp-In^™^293 cell lines. DOX-induced cell lines (18 h) were co-stained with primary antibodies to the FLAG epitope (red) and the ER-resident protein calnexin or KDEL (ER, green). Nuclei (blue) are shown in the merged image (**c**) Velocity sedimentation of endogenous Hrd1 and DOX-induced Hrd1-FH complexes on 10-40% sucrose gradients. Both samples and gradients were prepared in 1% LMNG. (d) Schematic of proteomic pipeline used to determine HCIPs for ER-resident E3s.

**Extended Data Fig. 2.**
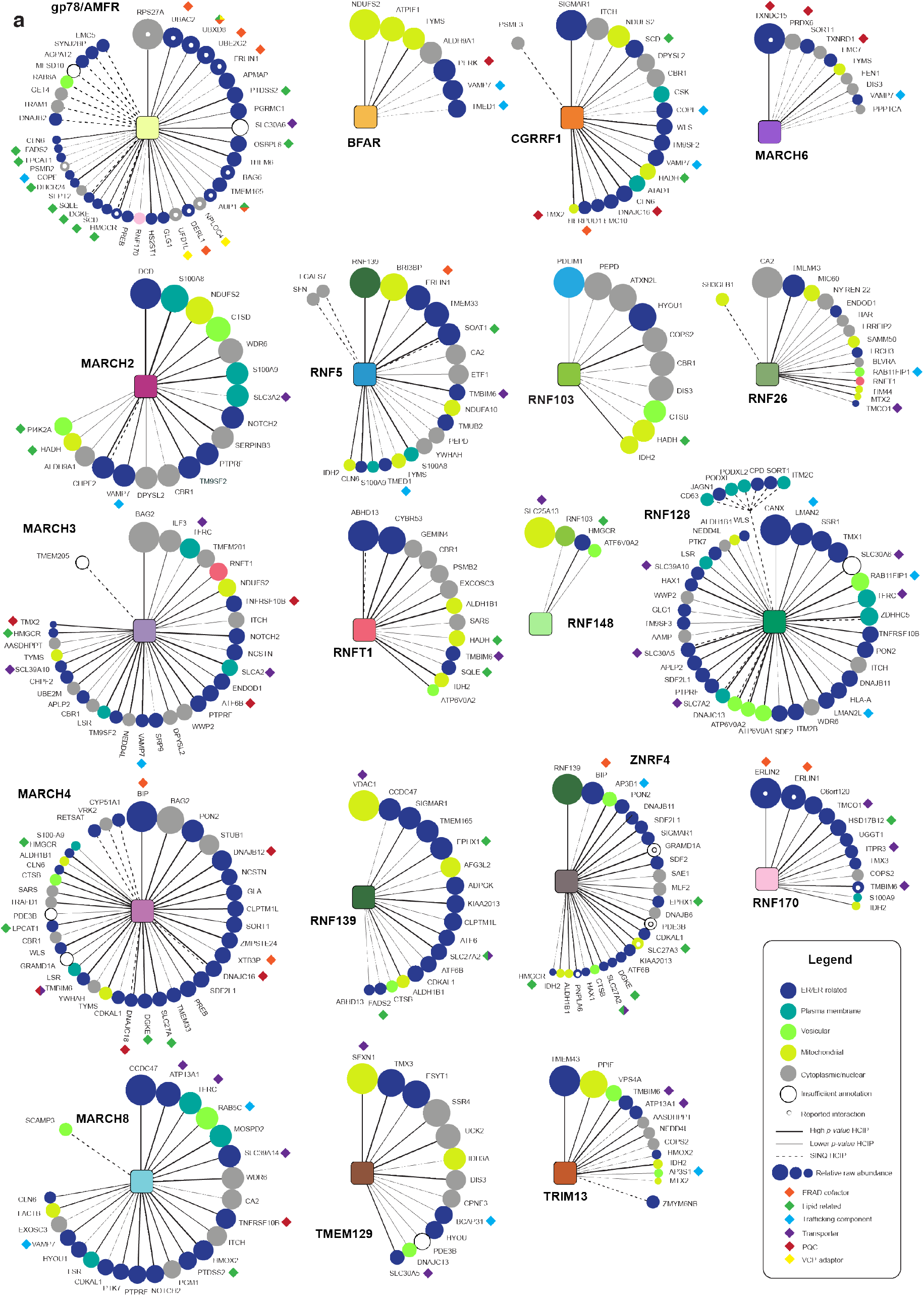
HCIP interaction network wheels for individual ER-E3s. (**a**) Protein-protein interaction wheels representing individual ER-resident E3 networks. Each HCIP is represented by a circle whose diameter reflects its raw abundance relative to the E3’s most abundant HCIP. Spoke/line thickness represents the p-value determined for each E3-HCIP interaction. Solid and dashed lines represent interactors identified by BCSG and SINQ analysis, respectively. Classifications of HCIP subcellular localisation are indicated by circle colour (described in legend), which have been manually curated using protein databases (e.g. UniProt) or when unavailable, assessed based on the predicted presence of organelle targeting features (e.g. signal sequence). HCIP function or related process, when known, is denoted by a proximal diamond (described in legend). HCIPs reported previously in BioGRID are indicated by a central white dot.

**Extended Data Fig. 3.**
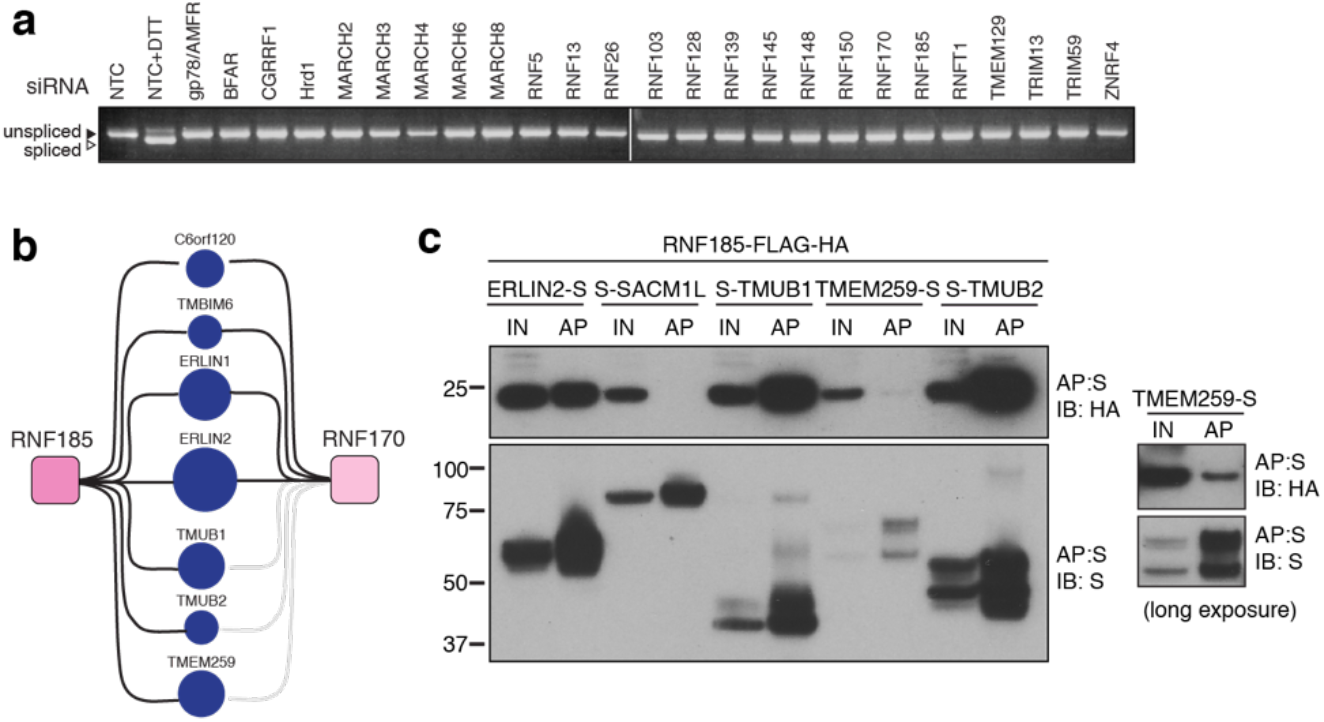
Validation of ER-resident E3 interactions. (**a**) Induction of ER stress as determined by splicing of *XBP1* in Flp-In^™^293 cells knocked down for individual ER-resident E3s by siRNAs. Splicing was validated by treatment of siNTC (non-targeting control)-transfected cells with DTT (5mM, 2 h). (**b**) Diagram representing shared HCIPs of RNF185 and RNF170. (**c**) Validation of RNF185 HCIPs. Transient expression of S-tagged HCIPs in Flp-In^™^293 cells stably expressing FH-RNF185. Complexes were affinity purified from LMNG lysates by S-protein agarose, separated by SDS-PAGE and the resulting western blots probed for RNF185 (anti-HA) and the HCIP (anti-S-tag). Because of weaker expression, TMEM259-S western blots are also presented in a longer exposure. Input (20%) and affinity purified (AP) material are shown.

**Extended Data Fig. 4.**
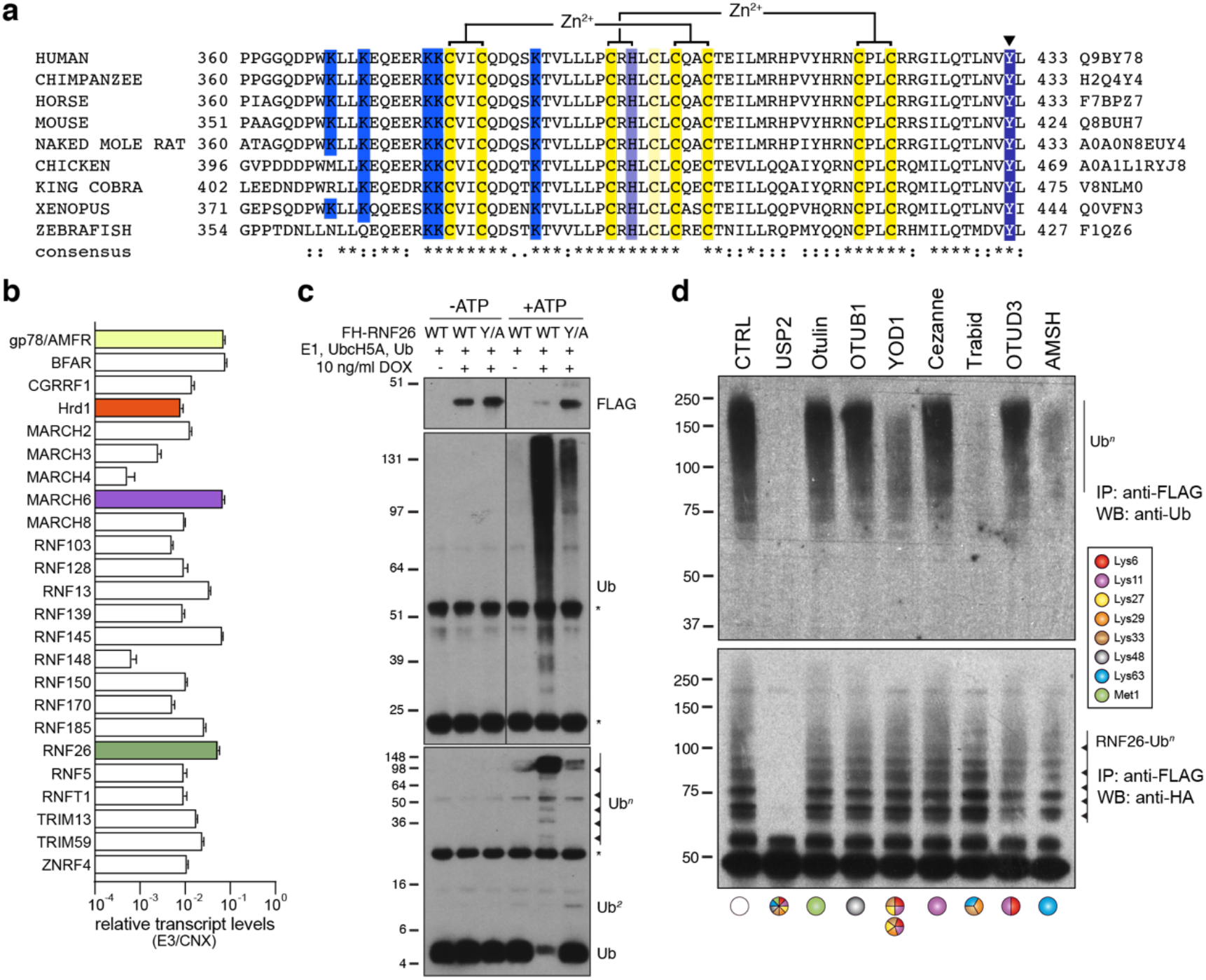
RNF26 ubiquitination. (**a**) Protein sequence alignment of the RING domain and C-terminus of RNF26 form different species. (**b**) E3 transcript abundance in Flp-In^™^293 cells determined by NanoString and normalised to calnexin levels. (**c**) FH-RNF26_WT_ or FH-RNF26_Y432A_ were isolated from their respective control or DOX-induced Flp-In^™^293 cell lines with anti-FLAG and combined with recombinant E1, UbcH5a, Ub, ± ATP to perform *in vitro* ubiquitination reactions. Resulting western blots were probed with antibodies against FLAG (RNF26) and Ub. Low (top panel) and high (bottom panel) percentage SDS-PAGE gels are presented to better resolve both poly- and mono-/di-ubiquitin forms, respectively. (**d**) Ub linkages present on FH-RNF26 as determined by UbiCREST analysis.

**Extended Data Fig. 5.**
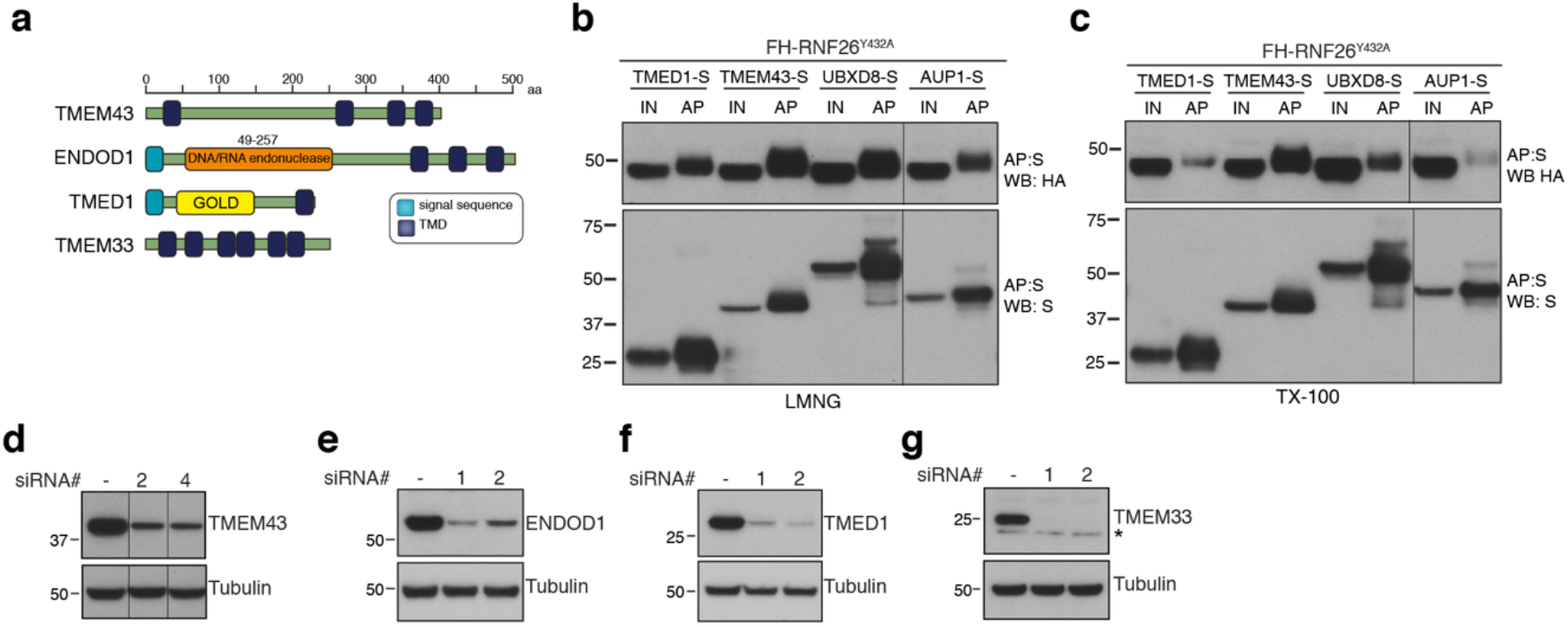
Validation of RNF26 interactions with HCIPs. (**a**) Domain organisation of human TMEM43, TMEM33, ENDOD1 and TMED1 proteins. Transient expression of S-tagged HCIPs in the FH-RNF26_Y432A_ Flp-In^™^293 cell line to validate robustness of RNF26-HCIP interactions. Cells solubilised in either 1% LMNG (**b**) or 1% TX-100 (**c**) yielded protein complexes affinity purified by S-protein agarose that were probed on western blots by antibodies against the S- and HA-tags. Input (IN, 20%) and affinity purified (AP) material are shown. (**d-g**) Validation of knockdown of TMEM43, TMEM33, ENDOD1 and TMED1 by two independent siRNAs. Western blots are probed using HCIP-specific antibodies and tubulin as a loading control.

**Extended Data Fig. 6.**
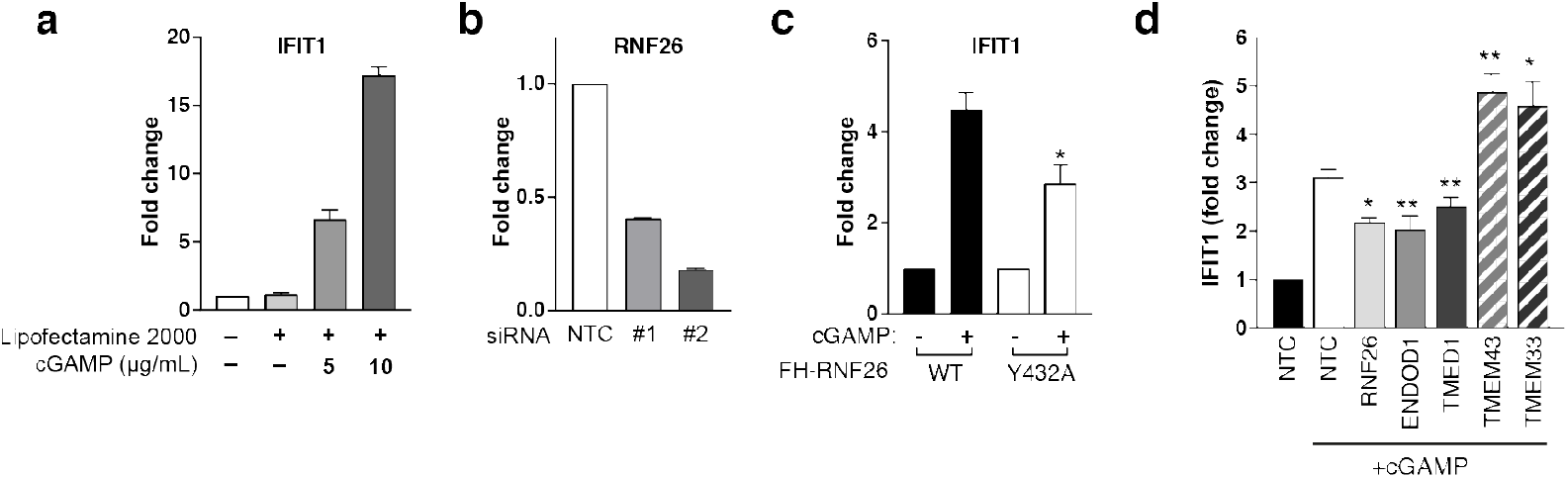
Modulation of the interferon response by RNF26 and its HCIPs. (**a**) Fold change in *IFIT1* transcription in response to cGAMP. Flp-In^™^293 cells were transfected with cGAMP (0, 5, 10 μg/ml) using Lipefectamine2000 (6 h) with *IFIT1* and *GAPDH* transcript levels measured by qRT-PCR (Taqman). Mean and S.E.M. from three biological replicates are presented (*n*=3). (**b**) Validation of RNF26 knockdown with two independent siRNAs by qPCR. Mean and S.E.M. from three biological repeats are presented. (**c**) Fold change in *IFIT1* transcription in response to cGAMP in FH-RNF26_WT_ and FH-RNF26_Y432A_ Flp-In^™^293 cell lines. Mean and S.E.M. from four biological replicates are presented. For all statistical analysis, \**P* < 0.05, \*\**P* < 0.01*,***P* < 0.001. Details of statistical analysis can be found in Extended Data Table 12. (**d**) As described in Fig. 6d for an independent set of siRNAs.

## Extended Data Tables (in corresponding Excel File)

**Extended Data Table 1:** ER-resident E3 features and positioning of FLAG-HA epitopes

**Extended Data Table 2:** Unique protein signatures of ER-resident E3 interactors identified by LC-MS/MS and BSCG analysis

**Extended Data Table 3:** Common contaminants removed from BSCG analysis

**Extended Data Table 4:** Unique ER-resident E3 interactors identified by SINQ analysis of LC-MS/MS dataset

**Extended Data Table 5:** High-confidence candidate interacting proteins (HCIPs) of ER-resident E3s

**Extended Data Table 6:** NanoString quantitative transcriptomics of ER-resident E3s in response to ER stress

**Extended Data Table 7:** Targets of UPR transcription factors among ER-resident E3 HCIPs

**Extended Data Table 8:** BSCG analysis of RNF26_Y432A_ compared to RNF26_WT_

**Extended Data Table 9:** MGC clones for ER-resident E3

**Extended Data Table 10:** MGC clones for HCIPs

**Extended Data Table 11:** siRNA sequences

**Extended Data Table 12:** Statistical analysis

## References

1. Valm, A. M. et al. Applying systems-level spectral imaging and analysis to reveal the organelle interactome. Nature 546, 162–167 (2017).

2. Kornmann, B. et al. An ER-Mitochondria Tethering Complex Revealed by a Synthetic Biology Screen. Science 325, 477–481 (2009).

3. Smith, J. J. & Aitchison, J. D. Peroxisomes take shape. Nat. Rev. Mol. Cell Biol. 14, 803–817 (2013).

4. van der Kant, R. & Neefjes, J. Small regulators, major consequences - Ca^2+^ and cholesterol at the endosome-ER interface. J. Cell Sci. 127, 929–938 (2014).

5. Hönscher, C. & Ungermann, C. A close-up view of membrane contact sites between the endoplasmic reticulum and the endolysosomal system: from yeast to man. Crit. Rev. Biochem. Mol. Biol. 49, 262–268 (2014).

6. Saheki, Y. & De Camilli, P. Endoplasmic Reticulum-Plasma Membrane Contact Sites. Annu. Rev. Biochem. 86, 659–684 (2017).

7. Wu, H., Carvalho, P. & Voeltz, G. K. Here, there, and everywhere: The importance of ER membrane contact sites. Science 361, eaan5835 (2018).

8. Komander, D. & Rape, M. The ubiquitin code. Annu. Rev. Biochem. 81, 203–229 (2012).

9. Li, W. et al. Genome-Wide and Functional Annotation of Human E3 Ubiquitin Ligases Identifies MULAN, a Mitochondrial E3 that Regulates the Organelle’s Dynamics and Signaling. PLoS ONE 3, e1487 (2008).

10. Zheng, N. & Shabek, N. Ubiquitin Ligases: Structure, Function, and Regulation. Annu. Rev. Biochem. 86, 129–157 (2017).

11. Foresti, O., Rodriguez-Vaello, V., Funaya, C. & Carvalho, P. Quality control of inner nuclear membrane proteins by the Asi complex. Science 346, 751–755 (2014).

12. Claessen, J. H. L., Kundrat, L. & Ploegh, H. L. Protein quality control in the ER: balancing the ubiquitin checkbook. Trends Cell Biol 22, 22–32 (2012).

13. Stewart, E. V. et al. Yeast SREBP cleavage activation requires the Golgi Dsc E3 ligase complex. Mol. Cell 42, 160–171 (2011).

14. Piper, R. C. & Lehner, P. J. Endosomal transport via ubiquitination. Trends Cell Biol 21, 647–655 (2011).

15. Schoebel, S. et al. Cryo-EM structure of the protein-conducting ERAD channel Hrd1 in complex with Hrd3. Nature 548, 352–355 (2017).

16. Vembar, S. S. & Brodsky, J. L. One step at a time: endoplasmic reticulum-associated degradation. Nat. Rev. Mol. Cell Biol. 9, 944–957 (2008).

17. Christianson, J. C. & Ye, Y. Cleaning up in the endoplasmic reticulum: ubiquitin in charge. Nat. Struct. Mol. Biol. 21, 325–335 (2014).

18. Hebert, D. N. & Molinari, M. In and out of the ER: protein folding, quality control, degradation, and related human diseases. Physiol Rev 87, 1377–1408 (2007).

19. Hampton, R. Y., Gardner, R. G. & Rine, J. Role of 26S proteasome and HRD genes in the degradation of 3-hydroxy-3-methylglutaryl-CoA reductase, an integral endoplasmic reticulum membrane protein. Mol. Biol. Cell 7, 2029–2044 (1996).

20. Carvalho, P., Goder, V. & Rapoport, T. A. Distinct ubiquitin-ligase complexes define convergent pathways for the degradation of ER proteins. Cell 126, 361–373 (2006).

21. Carvalho, P., Stanley, A. M. & Rapoport, T. A. Retrotranslocation of a misfolded luminal ER protein by the ubiquitin-ligase Hrd1p. Cell 143, 579–591 (2010).

22. Christianson, J. C., Shaler, T. A., Tyler, R. E. & Kopito, R. R. OS-9 and GRP94 deliver mutant alpha1-antitrypsin to the Hrd1-SEL1L ubiquitin ligase complex for ERAD. Nat. Cell Biol. 10, 272–282 (2008).

23. Bays, N. W., Gardner, R. G., Seelig, L. P., Joazeiro, C. A. & Hampton, R. Y. Hrd1p/Der3p is a membrane-anchored ubiquitin ligase required for ER-associated degradation. Nat. Cell Biol. 3, 24–29 (2001).

24. Christianson, J. C. et al. Defining human ERAD networks through an integrative mapping strategy. Nat. Cell Biol. 14, 93–105 (2012).

25. Stein, A., Ruggiano, A., Carvalho, P. & Rapoport, T. A. Key steps in ERAD of luminal ER proteins reconstituted with purified components. Cell 158, 1375–1388 (2014).

26. Hwang, J. et al. Characterization of protein complexes of the endoplasmic reticulum-associated degradation E3 ubiquitin ligase Hrd1. J. Biol. Chem. 292, 9104–9116 (2017).

27. Schulz, J. et al. Conserved cytoplasmic domains promote Hrd1 ubiquitin ligase complex formation for ER-associated degradation (ERAD). J. Cell Sci. 130, 3322–3335 (2017).

28. Ye, Y., Meyer, H. H. & Rapoport, T. A. The AAA ATPase Cdc48/p97 and its partners transport proteins from the ER into the cytosol. Nature 414, 652–656 (2001).

29. Ye, Y., Meyer, H. H. & Rapoport, T. A. Function of the p97-Ufd1-Npl4 complex in retrotranslocation from the ER to the cytosol: dual recognition of nonubiquitinated polypeptide segments and polyubiquitin chains. J. Cell Biol. 162, 71–84 (2003).

30. Ruggiano, A., Foresti, O. & Carvalho, P. Quality control: ER-associated degradation: protein quality control and beyond. J. Cell Biol. 204, 869–879 (2014).

31. Schwarz, D. S. & Blower, M. D. The endoplasmic reticulum: structure, function and response to cellular signaling. Cell. Mol. Life Sci. 73, 79–94 (2015).

32. Song, B.-L., Sever, N. & DeBose-Boyd, R. A. Gp78, a membrane-anchored ubiquitin ligase, associates with Insig-1 and couples sterol-regulated ubiquitination to degradation of HMG CoA reductase. Mol. Cell 19, 829–840 (2005).

33. Jo, Y., Lee, P. C. W., Sguigna, P. V. & DeBose-Boyd, R. A. Sterol-induced degradation of HMG CoA reductase depends on interplay of two Insigs and two ubiquitin ligases, gp78 and Trc8. Proc. Natl. Acad. Sci. U.S.A. 108, 20503–20508 (2011).

34. Jiang, L.-Y. et al. Ring finger protein 145 (RNF145) is a ubiquitin ligase for sterol-induced degradation of HMG-CoA reductase. J. Biol. Chem. 293, 4047–4055 (2018).

35. Menzies, S. A. et al. The sterol-responsive RNF145 E3 ubiquitin ligase mediates the degradation of HMG-CoA reductase together with gp78 and Hrd1. Elife 7, 24 (2018).

36. Foresti, O., Ruggiano, A., Hannibal-Bach, H. K., Ejsing, C. S. & Carvalho, P. Sterol homeostasis requires regulated degradation of squalene monooxygenase by the ubiquitin ligase Doa10/Teb4. Elife 2, e00953 (2013).

37. Zelcer, N. et al. The E3 ubiquitin ligase MARCH6 degrades squalene monooxygenase and affects 3-hydroxy-3-methyl-glutaryl coenzyme A reductase and the cholesterol synthesis pathway. Mol. Cell Biol. 34, 1262–1270 (2014).

38. Lu, J. P., Wang, Y., Sliter, D. A., Pearce, M. M. P. & Wojcikiewicz, R. J. H. RNF170 protein, an endoplasmic reticulum membrane ubiquitin ligase, mediates inositol 1,4,5-trisphosphate receptor ubiquitination and degradation. J. Biol. Chem. 286, 24426–24433 (2011).

39. Ishikawa, H. & Barber, G. N. STING is an endoplasmic reticulum adaptor that facilitates innate immune signalling. Nature 455, 674–678 (2008).

40. Cox, J. H., Yewdell, J. W., Eisenlohr, L. C., Johnson, P. R. & Bennink, J. R. Antigen presentation requires transport of MHC class I molecules from the endoplasmic reticulum. Science 247, 715–718 (1990).

41. Jongsma, M. L. M. et al. An ER-Associated Pathway Defines Endosomal Architecture for Controlled Cargo Transport. Cell 166, 152–166 (2016).

42. Zhao, Y., Zhang, T., Huo, H., Ye, Y. & Liu, Y. Lunapark Is a Component of a Ubiquitin Ligase Complex Localized to the Endoplasmic Reticulum Three-way Junctions. J. Biol. Chem. 291, 18252–18262 (2016).

43. Wu, Y. et al. Transmembrane E3 ligase RNF183 mediates ER stress-induced apoptosis by degrading Bcl-xL. Proc. Natl. Acad. Sci. U.S.A. 115, E2762–E2771 (2018).

44. Neutzner, A. et al. A systematic search for endoplasmic reticulum (ER) membrane-associated RING finger proteins identifies Nixin/ZNRF4 as a regulator of calnexin stability and ER homeostasis. J. Biol. Chem. 286, 8633–8643 (2011).

45. Maruyama, Y., Yamada, M., Takahashi, K. & Yamada, M. Ubiquitin ligase Kf-1 is involved in the endoplasmic reticulum-associated degradation pathway. Biochem. Biophys. Res. Commun. 374, 737–741 (2008).

46. van de Weijer, M. L. et al. A high-coverage shRNA screen identifies TMEM129 as an E3 ligase involved in ER-associated protein degradation. Nat. Commun. 5, – (2014).

47. Lilley, B. N. & Ploegh, H. L. Multiprotein complexes that link dislocation, ubiquitination, and extraction of misfolded proteins from the endoplasmic reticulum membrane. Proc. Natl. Acad. Sci. U.S.A. 102, 14296–14301 (2005).

48. Mueller, B., Klemm, E. J., Spooner, E., Claessen, J. H. & Ploegh, H. L. SEL1L nucleates a protein complex required for dislocation of misfolded glycoproteins. Proc. Natl. Acad. Sci. U.S.A. 105, 12325–12330 (2008).

49. Hosokawa, N., Wada, I., Okawa, K. & Nagata, K. Human XTP3-B forms an endoplasmic reticulum quality control scaffold with the HRD1-SEL1L ubiquitin ligase complex and BiP. J. Biol. Chem. 283, 20914–20924 (2008).

50. Keilhauer, E. C., Hein, M. Y. & Mann, M. Accurate Protein Complex Retrieval by Affinity Enrichment Mass Spectrometry (AE-MS) Rather than Affinity Purification Mass Spectrometry (AP-MS). Molecular & Cellular Proteomics 14, 120–135 (2015).

51. Sowa, M. E., Bennett, E. J., Gygi, S. P. & Harper, J. W. Defining the Human Deubiquitinating Enzyme Interaction Landscape. Cell 138, 389–403 (2009).

52. Trudgian, D. C. et al. Comparative evaluation of label-free SINQ normalized spectral index quantitation in the central proteomics facilities pipeline. Proteomics 11, 2790–2797 (2011).

53. Stark, C. et al. The BioGRID Interaction Database: 2011 update. Nucleic Acids Res. 39, D698–704 (2011).

54. Fang, S. et al. The tumor autocrine motility factor receptor, gp78, is a ubiquitin protein ligase implicated in degradation from the endoplasmic reticulum. Proc. Natl. Acad. Sci. U.S.A. 98, 14422–14427 (2001).

55. Travers, K. J. et al. Functional and genomic analyses reveal an essential coordination between the unfolded protein response and ER-associated degradation. Cell 101, 249–258 (2000).

56. Paton, A. W. et al. AB5 subtilase cytotoxin inactivates the endoplasmic reticulum chaperone BiP. Nature 443, 548–552 (2006).

57. Kaneko, M. et al. Genome-wide identification and gene expression profiling of ubiquitin ligases for endoplasmic reticulum protein degradation. Sci. Rep. 6, 30955 (2016).

58. Bergmann, T. J. et al. Chemical stresses fail to mimic the unfolded protein response resulting from luminal load with unfolded polypeptides. J. Biol. Chem. 293, jbc.RA117.001484–5612 (2018).

59. Adamson, B. et al. A Multiplexed Single-Cell CRISPR Screening Platform Enables Systematic Dissection of the Unfolded Protein Response. Cell 167, 1867–1882.e21 (2016).

60. Vitale, M. et al. Inadequate BiP availability defines endoplasmic reticulum stress. Elife 8, 74 (2019).

61. Ye, Y., Shibata, Y., Yun, C., Ron, D. & Rapoport, T. A. A membrane protein complex mediates retro-translocation from the ER lumen into the cytosol. Nature 429, 841–847 (2004).

62. Meyer, H., Bug, M. & Bremer, S. Emerging functions of the VCP/p97 AAA-ATPase in the ubiquitin system. Nat. Cell Biol. 14, 117–123 (2012).

63. Morreale, G., Conforti, L., Coadwell, J., Wilbrey, A. L. & Coleman, M. P. Evolutionary divergence of valosin-containing protein/cell division cycle protein 48 binding interactions among endoplasmic reticulum-associated degradation proteins. FEBS J. 276, 1208–1220 (2009).

64. Ballar, P., Shen, Y., Yang, H. & Fang, S. The role of a novel p97/valosin-containing protein-interacting motif of gp78 in endoplasmic reticulum-associated degradation. J. Biol. Chem. 281, 35359–35368 (2006).

65. Greenblatt, E. J., Olzmann, J. A. & Kopito, R. R. Derlin-1 is a rhomboid pseudoprotease required for the dislocation of mutant α-1 antitrypsin from the endoplasmic reticulum. Nat. Struct. Mol. Biol. 18, 1147–1152 (2011).

66. Ye, Y. et al. Recruitment of the p97 ATPase and ubiquitin ligases to the site of retrotranslocation at the endoplasmic reticulum membrane. Proc. Natl. Acad. Sci. U.S.A. 102, 14132–14138 (2005).

67. Fleig, L. et al. Ubiquitin-dependent intramembrane rhomboid protease promotes ERAD of membrane proteins. Mol. Cell 47, 558–569 (2012).

68. Khouri, El, E., Le Pavec, G., Toledano, M. B. & Delaunay-Moisan, A. RNF185 Is a Novel E3 Ligase of Endoplasmic Reticulum-associated Degradation (ERAD) That Targets Cystic Fibrosis Transmembrane Conductance Regulator (CFTR). J. Biol. Chem. 288, 31177–31191 (2013).

69. Hülsmann, J. et al. AP-SWATH Reveals Direct Involvement of VCP/p97 in Integrated Stress Response Signaling Through Facilitating CReP/PPP1R15B Degradation. Mol. Cell Proteomics 17, 1295–1307 (2018).

70. Buchberger, A., Schindelin, H. & Hänzelmann, P. Control of p97 function by cofactor binding. FEBS Lett 589, 2578–2589 (2015).

71. Jo, Y., Sguigna, P. V. & DeBose-Boyd, R. A. Membrane-associated ubiquitin ligase complex containing gp78 mediates sterol-accelerated degradation of 3-hydroxy-3-methylglutaryl-coenzyme A reductase. J. Biol. Chem. 286, 15022–15031 (2011).

72. Kny, M., Standera, S., Hartmann-Petersen, R., Kloetzel, P. M. & Seeger, M. Herp Regulates Hrd1-mediated Ubiquitylation in a Ubiquitin-like Domain-dependent Manner. J. Biol. Chem. 286, 5151–5156 (2011).

73. Morito, D. et al. Gp78 cooperates with RMA1 in endoplasmic reticulum-associated degradation of CFTRDeltaF508. Mol. Biol. Cell 19, 1328–1336 (2008).

74. Zhang, T., Xu, Y., Liu, Y. & Ye, Y. gp78 functions downstream of Hrd1 to promote degradation of misfolded proteins of the endoplasmic reticulum. Mol. Biol. Cell 26, 4438–4450 (2015).

75. Pearce, M. M. P., Wang, Y., Kelley, G. G. & Wojcikiewicz, R. J. H. SPFH2 mediates the endoplasmic reticulum-associated degradation of inositol 1,4,5-trisphosphate receptors and other substrates in mammalian cells. J. Biol. Chem. 282, 20104–20115 (2007).

76. Pearce, M. M. P., Wormer, D. B., Wilkens, S. & Wojcikiewicz, R. J. H. An Endoplasmic Reticulum (ER) Membrane Complex Composed of SPFH1 and SPFH2 Mediates the ER-associated Degradation of Inositol 1,4,5-Trisphosphate Receptors. J. Biol. Chem. 284, 10433–10445 (2009).

77. Bultynck, G., Kiviluoto, S. & Methner, A. Bax inhibitor-1 is likely a pH-sensitive calcium leak channel, not a H+/Ca2+ exchanger. Sci. Signal. 7, pe22–pe22 (2014).

78. Rojas-Rivera, D. et al. TMBIM3/GRINA is a novel unfolded protein response (UPR) target gene that controls apoptosis through the modulation of ER calcium homeostasis. Cell Death Differ 19, 1013–1026 (2012).

79. Yang, B. et al. The critical role of membralin in postnatal motor neuron survival and disease. Elife 4, 510 (2015).

80. Zhu, B. et al. ER-associated degradation regulates Alzheimer’s amyloid pathology and memory function by modulating γ-secretase activity. Nat. Commun. 8, 1472 (2017).

81. Wang, Q.-C. et al. TMCO1 Is an ER Ca2+ Load-Activated Ca2+ Channel. Cell 165, 1454–1466 (2016).

82. Kang, R. et al. Neural palmitoyl-proteomics reveals dynamic synaptic palmitoylation. Nature 456, 904–909 (2008).

83. Martin, B. R. & Cravatt, B. F. Large-scale profiling of protein palmitoylation in mammalian cells. Nat. Methods 6, 135–138 (2009).

84. Breusegem, S. Y. & Seaman, M. N. J. Genome-wide RNAi Screen Reveals a Role for Multipass Membrane Proteins in Endosome-to-Golgi Retrieval. Cell Rep 9, 1931–1945 (2014).

85. Anandasabapathy, N. et al. GRAIL: an E3 ubiquitin ligase that inhibits cytokine gene transcription is expressed in anergic CD4+ T cells. Immunity 18, 535–547 (2003).

86. Qin, Y. et al. RNF26 temporally regulates virus-triggered type I interferon induction by two distinct mechanisms. PLoS Pathog. 10, e1004358 (2014).

87. Liew, C. W., Sun, H., Hunter, T. & Day, C. L. RING domain dimerization is essential for RNF4 function. Biochem J 431, 23–29 (2010).

88. Plechanovová, A., Jaffray, E. G., Tatham, M. H., Naismith, J. H. & Hay, R. T. Structure of a RING E3 ligase and ubiquitin-loaded E2 primed for catalysis. Nature 489, 115–120 (2012).

89. Gyrd-Hansen, M. et al. IAPs contain an evolutionarily conserved ubiquitin-binding domain that regulates NF-kappaB as well as cell survival and oncogenesis. Nat. Cell Biol. 10, 1309–1317 (2008).

90. Linke, K. et al. Structure of the MDM2/MDMX RING domain heterodimer reveals dimerization is required for their ubiquitylation in trans. Cell Death Differ 15, 841–848 (2008).

91. Plechanovová, A. et al. Mechanism of ubiquitylation by dimeric RING ligase RNF4. Nat. Struct. Mol. Biol. 18, 1052–1059 (2011).

92. Bengtsson, L. & Otto, H. LUMA interacts with emerin and influences its distribution at the inner nuclear membrane. J. Cell Sci. 121, 536–548 (2008).

93. Dreger, M., Bengtsson, L., Schöneberg, T., Otto, H. & Hucho, F. Nuclear envelope proteomics: Novel integral membrane proteins of the inner nuclear membrane. Proc. Natl. Acad. Sci. U.S.A. 98, 11943–11948 (2001).

94. Jenne, N., Frey, K., Brügger, B. & Wieland, F. T. Oligomeric state and stoichiometry of p24 proteins in the early secretory pathway. J. Biol. Chem. 277, 46504–46511 (2002).

95. Urade, T., Yamamoto, Y., Zhang, X., Ku, Y. & Sakisaka, T. Identification and characterization of TMEM33 as a reticulon-binding protein. Kobe J Med Sci 60, E57–65 (2014).

96. Tyler, R. E. et al. Unassembled CD147 is an endogenous endoplasmic reticulum-associated degradation substrate. Mol. Biol. Cell 23, 4668–4678 (2012).

97. Bridgeman, A. et al. Viruses transfer the antiviral second messenger cGAMP between cells. Science 349, 1228–1232 (2015).

98. Shang, G., Zhang, C., Chen, Z. J., Bai, X.-C. & Zhang, X. Cryo-EM structures of STING reveal its mechanism of activation by cyclic GMP–AMP. Nature 339, 1 (2019).

99. Ishikawa, H., Ma, Z. & Barber, G. N. STING regulates intracellular DNA-mediated, type I interferon-dependent innate immunity. Nature 461, 788–792 (2009).

100. Dobbs, N. et al. STING Activation by Translocation from the ER Is Associated with Infection and Autoinflammatory Disease. Cell Host Microbe 18, 157–168 (2015).

101. Prabakaran, T. et al. Attenuation of cGAS-STING signaling is mediated by a p62/SQSTM1-dependent autophagy pathway activated by TBK1. EMBO J 37, e97858 (2018).

102. Zhong, B. et al. The Ubiquitin Ligase RNF5 Regulates Antiviral Responses by Mediating Degradation of the Adaptor Protein MITA. Immunity 30, 397–407 (2009).

103. Wang, Q. et al. The E3 Ubiquitin Ligase AMFR and INSIG1 Bridge the Activation of TBK1 Kinase by Modifying the Adaptor STING. Immunity 41, 919–933 (2014).

104. Bennett, E. J., Rush, J., Gygi, S. P. & Harper, J. W. Dynamics of cullin-RING ubiquitin ligase network revealed by systematic quantitative proteomics. Cell 143, 951–965 (2010).

105. Glaeser, K. et al. ERAD-dependent control of the Wnt secretory factor Evi. EMBO J 37, e97311 (2018).

106. Prabhu, A. V., Luu, W., Sharpe, L. J. & Brown, A. J. Cholesterol-mediated Degradation of 7-Dehydrocholesterol Reductase Switches the Balance from Cholesterol to Vitamin D Synthesis. J. Biol. Chem. 291, 8363–8373 (2016).

107. Huang, E. Y. et al. A VCP inhibitor substrate trapping approach (VISTA) enables proteomic profiling of endogenous ERAD substrates. Mol. Biol. Cell 29, 1021–1030 (2018).

108. Küch, E.-M. et al. Differentially localized acyl-CoA synthetase 4 isoenzymes mediate the metabolic channeling of fatty acids towards phosphatidylinositol. Biochim Biophys Acta 1841, 227–239 (2014).

109. McShane, E. et al. Kinetic Analysis of Protein Stability Reveals Age-Dependent Degradation. Cell 167, 803–815.e21 (2016).

110. Locke, M., Toth, J. I. & Petroski, M. D. K11- and K48-Linked Ubiquitin Chains Interact with p97 during Endoplasmic Reticulum-Associated Degradation. Biochem J 459, 205–216 (2014).

111. Xu, P. et al. Quantitative proteomics reveals the function of unconventional ubiquitin chains in proteasomal degradation. Cell 137, 133–145 (2009).

112. Kristariyanto, Y. A. et al. K29-Selective Ubiquitin Binding Domain Reveals Structural Basis of Specificity and Heterotypic Nature of K29 Polyubiquitin. Molecular Cell 58, 83–94 (2015).

113. Leto, D. E. et al. Parallel genome-wide CRISPR analysis identifies a role for heterotypic ubiquitin chains in ER-associated degradation. bioRxiv 349407 (2018). doi:10.1101/349407

114. Das, R. et al. Allosteric activation of E2-RING finger-mediated ubiquitylation by a structurally defined specific E2-binding region of gp78. Mol. Cell 34, 674–685 (2009).

115. van Wijk, S. J. L. et al. A comprehensive framework of E2-RING E3 interactions of the human ubiquitin-proteasome system. Mol. Syst. Biol. 5, 295 (2009).

116. Markson, G. et al. Analysis of the human E2 ubiquitin conjugating enzyme protein interaction network. Genome Res. 19, 1905–1911 (2009).

117. Shoulders, M. D. et al. Stress-independent activation of XBP1s and/or ATF6 reveals three functionally diverse ER proteostasis environments. Cell Rep 3, 1279–1292 (2013).

118. Yagishita, N. et al. Essential role of synoviolin in embryogenesis. J. Biol. Chem. 280, 7909–7916 (2005).

119. Vasic, V. et al. Hrd1 forms the retrotranslocation pore regulated by auto-ubiquitination and binding of misfolded proteins. Nat. Cell Biol. 21, 1–8 (2020).

120. van den Boomen, D. J. H. et al. TMEM129 is a Derlin-1 associated ERAD E3 ligase essential for virus-induced degradation of MHC-I. Proc. Natl. Acad. Sci. U.S.A. 111, 11425–11430 (2014).

121. Lilley, B. N., Spooner, E., Tortorella, D. & Ploegh, H. L. Signal peptide peptidase is required for dislocation from the endoplasmic reticulum. Nature 441, 894–897 (2006).

122. Wickner, W. & Schekman, R. Protein Translocation Across Biological Membranes. Science 310, 1452–1456 (2005).

123. Cai, X., Chiu, Y.-H. & Chen, Z. J. The cGAS-cGAMP-STING pathway of cytosolic DNA sensing and signaling. Molecular Cell 54, 289–296 (2014).

124. Zhang, D., Vjestica, A. & Oliferenko, S. The cortical ER network limits the permissive zone for actomyosin ring assembly. Curr Biol 20, 1029–1034 (2010).

125. Zhang, D. & Oliferenko, S. Tts1, the fission yeast homologue of the TMEM33 family, functions in NE remodeling during mitosis. Mol. Biol. Cell 25, 2970–2983 (2014).

126. Merner, N. D. et al. Arrhythmogenic right ventricular cardiomyopathy type 5 is a fully penetrant, lethal arrhythmic disorder caused by a missense mutation in the TMEM43 gene. Am. J. Hum. Genet. 82, 809–821 (2008).

127. Liang, W. C. et al. TMEM43 mutations in Emery-Dreifuss muscular dystrophy-related myopathy. Ann. Neurol. 69, 1005–1013 (2011).

128. Jiang, C. et al. TMEM43/LUMA is a key signaling component mediating EGFR-induced NF-κB activation and tumor progression. Oncogene 36, 2813–2823 (2017).

129. Jin, L. et al. Ubiquitin-dependent regulation of COPII coat size and function. Nature 482, 495–500 (2012).

130. Lyu, Z.-Z., Zhao, B.-B., Koiwai, K., Hirono, I. & Kondo, H. Identification of endonuclease domain-containing 1 gene in Japanese flounder Paralichthys olivaceus. Fish Shellfish Immunol. 50, 43–49 (2016).

131. Hospenthal, M. K., Mevissen, T. E. T. & Komander, D. Deubiquitinase-based analysis of ubiquitin chain architecture using Ubiquitin Chain Restriction (UbiCRest). Nat. Protoc. 10, 349–361 (2015).

132. Finn, R. D. et al. The Pfam protein families database: towards a more sustainable future. Nucleic Acids Res. 44, D279–85 (2016).

133. Tsirigos, K. D., Peters, C., Shu, N., Käll, L. & Elofsson, A. The TOPCONS web server for consensus prediction of membrane protein topology and signal peptides. Nucleic Acids Res. 43, W401–7 (2015).

